# A Molecular Mechanism for Turning off IRE1α Signaling During Endoplasmic Reticulum Stress

**DOI:** 10.1101/2020.04.03.024356

**Authors:** Xia Li, Sha Sun, Suhila Appathurai, Arunkumar Sundaram, Rachel Plumb, Malaiyalam Mariappan

## Abstract

Misfolded proteins in the endoplasmic reticulum (ER) activate IRE1α endoribonuclease in mammalian cells, which mediates XBP1 mRNA splicing to produce an active transcription factor. This promotes the expression of specific genes to alleviate ER stress and thereby attenuating IRE1α. Although sustained activation of IRE1α is linked to human diseases, it is not clear how IRE1α is attenuated during ER stress. Here, we identify that Sec63 is a subunit of the previously identified IRE1α/Sec61 translocon complex. We find that Sec63 recruits and activates BiP ATPase through its luminal J-domain to bind onto IRE1α. This leads to inhibition of higher-order oligomerization and attenuation of IRE1α RNase activity during prolonged ER stress. In Sec63 deficient cells, IRE1α remains activated for a long time despite the presence of excess BiP in the ER. Thus, our data suggest that the Sec61 translocon bridges IRE1α with Sec63/BiP to regulate the dynamics of IRE1α signaling in cells.

## Introduction

Secretory and membrane proteins are initially synthesized and folded in the endoplasmic reticulum (ER). The majority of these nascent proteins are delivered to the Sec61 translocon in the ER membrane by the co-translational protein targeting pathway (Rapoport, 2007; Shao and Hegde, 2011). The Sec61 translocon facilitates the translocation and insertion of newly synthesized secretory and membrane proteins. Immediately after entering the ER, they are folded and assembled with the help of glycosylation, chaperones, and folding enzymes in the ER (van Anken and Braakman, 2005). However, the ER capacity to fold newly synthesized proteins is often challenged by several conditions, including a sudden increase in incoming protein load, expression of aberrant proteins, and environmental stress. Under such conditions, terminally misfolded and unassembled proteins are recognized by the ER-associated degradation (ERAD) pathway for proteasomal degradation (Brodsky, 2012). When misfolded proteins overwhelm the ERAD capacity, they accumulate in the ER and thereby causing ER stress, which in turn triggers a signaling network called the unfolded protein response (UPR) (Walter and Ron, 2011). The UPR restores ER homeostasis by both reducing incoming protein load as well as increasing the protein folding capacity of the ER. If ER stress is unmitigated, the UPR has been shown to initiate apoptosis to eliminate non-functional cells (Hetz, 2012). The UPR-mediated life-and-death decision is implicated in several human diseases, including diabetes, cancer, and neurodegeneration (Hetz et al., 2020; Wang and Kaufman, 2016).

Three major transmembrane ER stress sensor proteins are localized in the mammalian ER, namely IRE1α, PERK, and ATF6α (Walter and Ron, 2011). IRE1α is a conserved transmembrane kinase/endonuclease, which is activated by self-oligomerization and trans-autophosphorylation during ER stress conditions (Cox et al., 1993; Mori et al., 1993). Once activated, IRE1α mediates nonconventional splicing of XBP1 mRNA (Calfon et al., 2002; Yoshida et al., 2001), which is recruited to the Sec61 translocon through its ribosome nascent chain (Plumb et al., 2015; Yanagitani et al., 2011). The cleaved fragments of XBP1 mRNA are subsequently ligated by the RtcB tRNA ligase (Jurkin et al., 2014; Kosmaczewski et al., 2014; Lu et al., 2014) with its co-factor archease (Poothong et al., 2017). The spliced XBP1 mRNA is translated into a functional transcription factor, XBP1s, which induces the expression of chaperones, quality control factors, and protein translocation components (Lee et al., 2003) (Lee et al., 2003). IRE1α can also promiscuously cleave many ER-localized mRNAs through the regulated Ire1-dependent decay (RIDD) pathway, which is implicated in incoming protein load to the ER as well as repositioning lysosomes during ER stress (Bae et al., 2019; Han et al., 2009; Hollien and Weissman, 2006). PERK is a transmembrane kinase and is responsible for phosphorylating the α subunit of eIF2 during ER stress, which causes global inhibition of translation in cells, thus alleviating the burden of protein misfolding in the ER (Harding et al., 1999; Sood et al., 2000). ATF6α is an ER-localized transcription factor and is translocated to Golgi upon ER stress where it is cleaved by intramembrane proteases (Haze et al., 1999; Ye et al., 2000). This causes the release of the cytosolic transcription domain into the cytosol and to the nucleus where it upregulates genes encoding ER chaperones and quality control factors to restore ER homeostasis (Lee et al., 2003; Shoulders et al., 2013).

The activity of all three UPR sensors is tightly regulated both under homeostatic and ER stress conditions, but the underlying mechanisms are unclear. In particular, it is important to understand the regulation of IRE1α activity since sustained activation of IRE1α is implicated in human diseases including type 2 diabetes (Ghosh et al., 2014; Hetz, 2012; Lin et al., 2007). On the other hand, hyperactivated IRE1α can produce an excess of XBP1s transcription factor, which can be beneficial for tumor cell growth in a hostile environment (Cubillos-Ruiz et al., 2017). Recent studies have identified many IRE1α interacting proteins that have been shown to regulate IRE1α activation and inactivation during ER stress (Eletto et al., 2014; Sepulveda et al., 2018; Sundaram et al., 2017). One of the key factors that regulates IRE1α activity is the luminal Hsp70 like chaperone BiP ATPase (Amin-Wetzel et al., 2017; Bertolotti et al., 2000; Carrara et al., 2015; Oikawa et al., 2009; Okamura et al., 2000; Pincus et al., 2010). However, it is unclear how the luminal protein BiP is efficiently recruited to the membrane-localized IRE1α in cells. Our previous studies have shown that IRE1α interaction with the Sec61 translocon is essential to suppress its oligomerization and RNase activity in cells (Sundaram et al., 2017). The molecular mechanism by which the Sec61 translocon limits IRE1α activity is unclear. In this study, we found that the Sec61 translocon bridges the interaction between IRE1α and Sec63. The J-domain of Sec63 is responsible for recruiting and activating the luminal BiP ATPase to bind onto IRE1α, thus suppressing IRE1α higher-order oligomerization and RNase activity during prolonged ER stress conditions.

## Results

### Sec63 is a subunit of the IRE1α/Sec61 translocon complex

We hypothesized that a Sec61 translocon associated protein may inhibit IRE1α oligomerization and RNase activity. This would explain why the Sec61 translocon interaction defective IRE1α mutants are resistant to attenuation during persistent ER stress (Sundaram et al., 2017). We therefore looked back at our previous results on IRE1α interacting proteins (Plumb et al., 2015). In addition to the Sec61 translocon, Sec63 is also enriched in the affinity-purified IRE1α sample from HEK293 cells. Sec63 is a conserved translocon interacting membrane protein, which contains a luminal J domain that activates the ATPase activity of BiP. Earlier studies have shown that Sec63 is required for both co-and post-translational protein translocation into the ER (Brodsky et al., 1995; Deshaies et al., 1991; Linxweiler et al., 2017; Meyer et al., 2000; Panzner et al., 1995). To confirm Sec63 is a part of the IRE1α/Sec61 translocon complex, we immunoprecipitated IRE1α from HEK293 cells expressing HA-tagged IRE1α. We also used several IRE1α variants carrying mutations within the 10 amino acid region located at the luminal juxtamembrane position that is critical for the interaction with the Sec61 translocon (Figure 1A) (Plumb et al., 2015). We used the lysis buffer containing digitonin detergent, as it preserves the interaction between IRE1α and the Sec61 translocon (Plumb et al., 2015). Indeed, IRE1α associated with Sec63 along with the Sec61 translocon (Figure 1B). Interestingly, IRE1α did not coimmunoprecipitate with Sec62, which is known to form a complex with Sec61/Sec63 (Panzner et al., 1995). IRE1α mutants that weakly interact with the Sec61 translocon also showed a significantly reduced interaction with Sec63 (Figure 1B). In addition to the luminal juxtamembrane region, the transmembrane domain (TMD) of IRE1α is also important for the interaction with Sec61/Sec63 since replacing IRE1α TMD with the TMD from Calnexin abolished the interaction with the translocon complex (Figures 1A and 1B).

**Figure 1.**
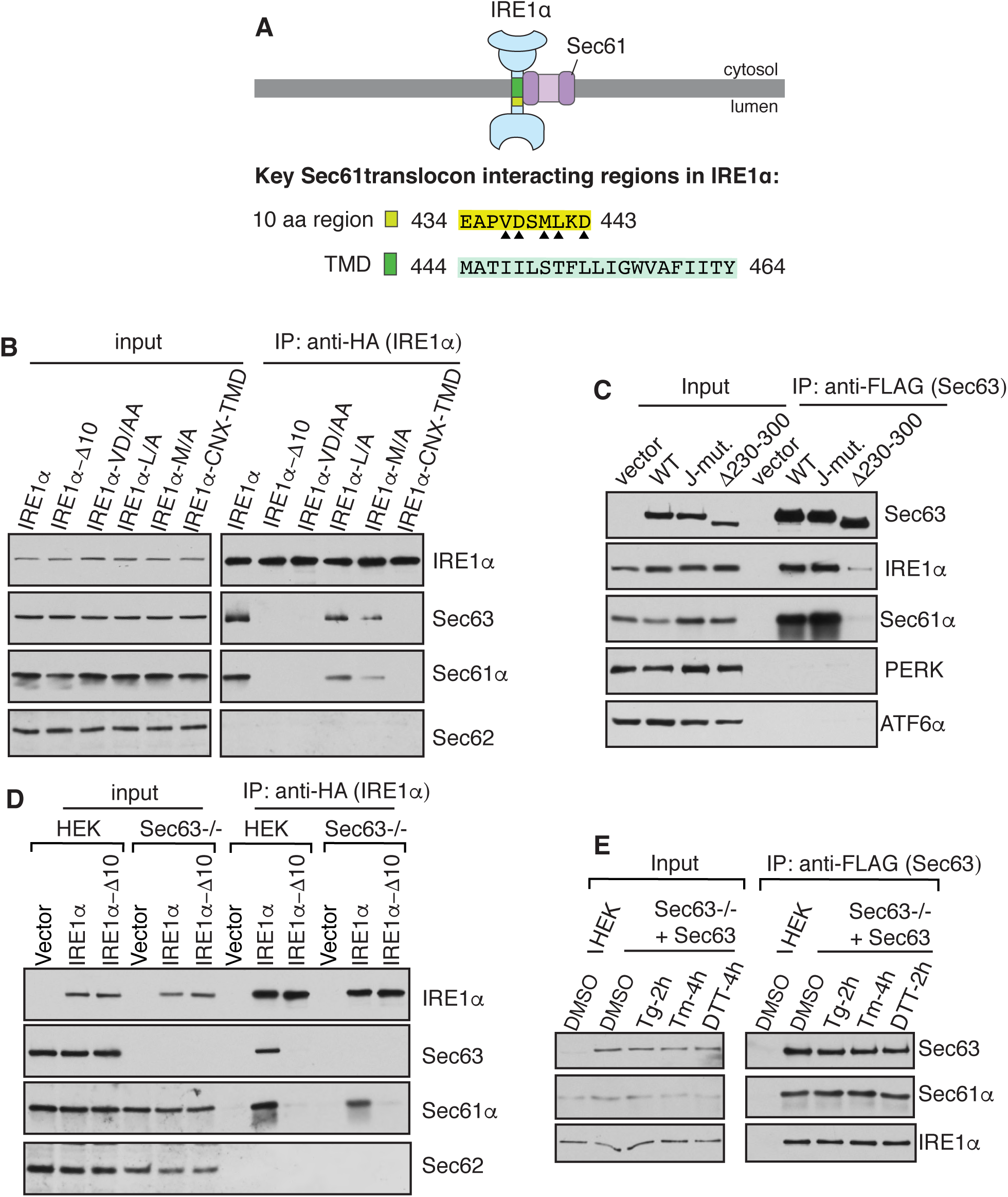
The Sec61 translocon bridges the interaction between IRE1α and Sec63. (A) A diagram showing the Sec61 translocon interacting regions in IRE1α. The 10 amino acid (aa) region (yellow) located at the luminal juxtamembrane position is required for the interaction with the Sec61 translocon (Plumb et al., 2015). Triangle depicts amino acid residues within the 10 aa region that are important for the interaction with the Sec61 translocon. In this study, we identified the transmembrane domain (green) of IRE1α is also required for the interaction with the Sec61 translocon. (B) The cell lysates expressing the indicated versions of HA-tagged IRE1α were immunoprecipitated using an anti-HA antibody and analyzed by immunoblotting. IRE1α Δ10 lacks amino acids 434 to 443 in human IRE1α. IRE1α VD/AA, L/A, and M/A are mutations within the 10-amino acid region shown in (A). The TMD of IRE1α is replaced with the TMD of calnexin in the IRE1α-CNX-TMD construct. (C) The cell lysates from FLAG-tagged wild type (WT) Sec63, the J-domain mutant, or Δ230-300 expressing cells were immunoprecipitated using an anti-FLAG antibody and immunoblotted for the indicated antigens. (D) HEK293 or HEK293 Sec63-/- cells were transfected with either wild type IRE1α-HA or IRE1α Δ10-HA and immunoprecipitated and analyzed as in (B). (E) Sec63-/- cells complemented with Sec63-FLAG were either treated with DMSO, 5 μg/ml thapsigargin (Tg) for 2h, 5 μg/ml tunicamycin (Tm) for 4h, or 4 mM DTT for 2h. Sec63 was immunoprecipitated from these cell lysates and immunoblotted for the indicated antigens. See also **Figure S1**.

The aforementioned data are compatible with two possible models. First, Sec63 interacts with IRE1α through the Sec61 translocon. Second, the regions in IRE1α (Figure 1A) that contribute to the interaction with Sec61 translocon may also be required for the interaction with Sec63. To differentiate between these two models, we sought to identify Sec63 mutants that disrupt the interaction with the Sec61 translocon. Sec63 mutants that poorly interacted with the Sec61 translocon also showed less interaction with the endogenous IRE1α, thus supporting the first model (Figures 1C, S1A, and S1B). Sec61/Sec63 selectively interacted with the IRE1α branch of the UPR since they did not appreciably interact with either PERK or ATF6α (Figure 1C). The first model is further corroborated by the observation that IRE1α interaction with the Sec61 translocon was less disrupted in Sec63-/- cells compared to wild type cells, suggesting that IRE1α can form a complex with the Sec61 translocon independent of Sec63 (Figure 1D).

Furthermore, we examined whether the interaction between Sec63 and IRE1α is preserved during ER stress conditions. To test this, we immunoprecipitated Sec63 from Sec63- /- cells complemented with Sec63-FLAG that were treated without or with an ER stress inducer, thapsigargin (Tg), tunicamycin (Tm), or dithiothreitol (DTT). Sec61/Sec63 interaction with the endogenous IRE1α was mostly undisrupted during ER stress conditions (Figure 1E and Figure S1C). Taken together, our data suggest that IRE1α interacts with Sec63 via the Sec61 translocon and the interaction is stable during ER stress.

### Sec63 suppresses the formation of higher-order oligomers of IRE1α during ER stress

It is known that IRE1α forms higher-order oligomers or clusters in cells upon ER stress, which correlate with IRE1α RNase activity (Li et al., 2010). We have previously shown that IRE1α interaction with the Sec61 translocon is crucial for limiting the formation of IRE1α clusters in cells during ER stress (Sundaram et al., 2017). Since Sec63 is also a subunit of the IRE1α/Sec61 translocon complex (Figure 1A), we hypothesized that Sec63 may be responsible for limiting IRE1α oligomerization during ER stress. To test this idea, we performed siRNA mediated knockdown of Sec63 in cells and monitored IRE1α clustering by confocal immunofluorescence after treating cells with the ER stress inducer Tg. IRE1α was localized to the ER without clustering under homeostatic conditions, while a small number of cells exhibited IRE1α clusters upon ER stress in control siRNA treated cells (Figures 2A, 2B, and 2C). By contrast, the number of IRE1α clusters was significantly increased in Sec63 depleted cells treated with Tg (Figures 2A, 2B, and 2C). An alternative explanation for IRE1α clustering in Sec63 depleted cells is that it may be caused by defects in protein translocation in these cells.

**Figure 2.**
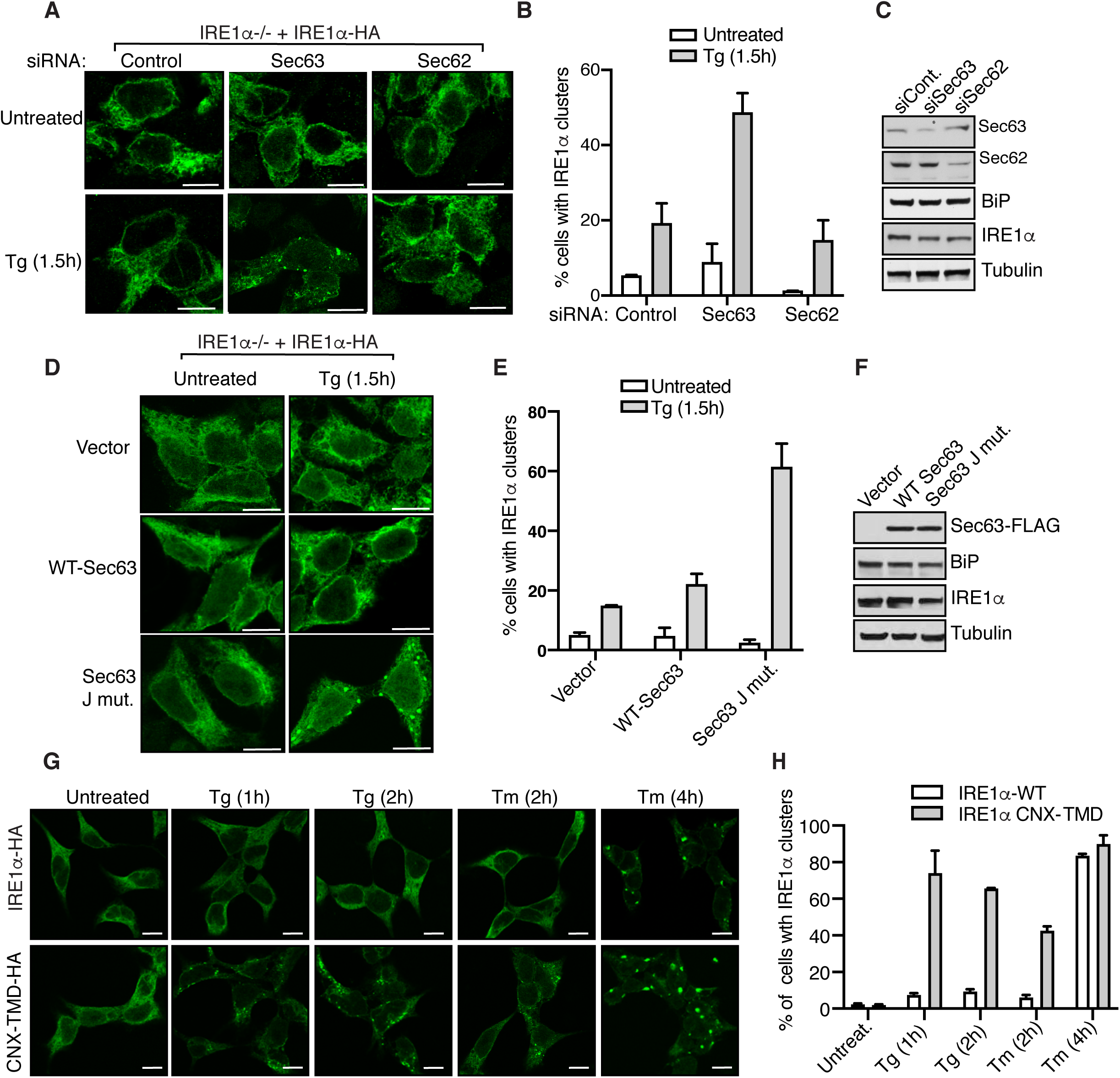
Sec63 inhibits IRE1α clustering during ER stress in cells. (A) HEK293 IRE1α-/- cells complemented with IRE1α-HA were transfected with either control, Sec63, or Sec62 siRNA. The expression of IRE1α-HA was induced with 5 ng/ml doxycycline. After 30h of transfection, the cells were either left untreated or treated with 5 μg/ml Tg for 1.5 h and processed for immunostaining with anti-HA antibodies for IRE1α. Scale bars are 10 μm. (B) Quantification results of the number of cells with IRE1α clusters in (A). Error bars represent the standard error of the mean (SEM) from two independent experiments. (C) HEK293 IRE1α-/- cells were treated with the indicated siRNAs as in (A) but analyzed by immunoblotting for the indicated antigens. (D) HEK293 IRE1α-/- cells expressing IRE1α-HA were transfected with either empty vector, wild type (WT) Sec63 or the J-domain mutant of Sec63. The expression of IRE1α-HA was induced with 5 ng/ml doxycycline. After 24 h of transfection, the cells were either left untreated or treated with 5 μg/ml Tg for 1.5 h and subsequently processed for immunostaining with anti-HA antibodies for IRE1α. (E) Quantification results of the number of cells with IRE1α clusters in (D). Error bars represent the standard error of the mean (SEM) from two independent experiments. (F) The cells were transfected as in (D) but analyzed by immunoblotting for the indicated antigens. (G) HEK293 IRE1α-/- cells expressing either IRE1α-HA or IRE1α-CNX-TMD-HA were induced with 5 ng/ml doxycycline and treated with either Tg or Tm for the indicated time points. The cells were then processed for immunostaining as in (A). (H) Quantification results of the number of cells with IRE1α clusters in (G). Error bars represent the standard error of the mean (SEM) from two independent experiments.

To rule out this possibility, we performed siRNA mediated knockdown of Sec62 and monitored IRE1α clusters in cells. Sec62 is also a translocon associated protein and is involved in protein translocon into the ER (Linxweiler et al., 2017). Unlike Sec63, transient depletion of Sec62 did not significantly increase IRE1α clusters upon ER stress compared to control siRNA treated cells (Figure 2A, 2B, and 2C). We next investigated whether the J-domain of Sec63 is required for suppressing IRE1α clustering in cells. The cells expressing the Sec63 J-domain mutant (HPD/AAA), which is deficient in activating the ATPase activity of BiP, showed more IRE1α clusters upon ER stress compared to cells expressing wild type Sec63 (Figures 2D, 2E, and 2F).

To further strengthen our conclusion that Sec63 inhibits IRE1α clustering, we monitored IRE1α clusters in cells expressing IRE1α CNX-TMD mutant, which we identified in this study as a Sec61/Sec63 interaction defective mutant (Figure 1B). Indeed, IRE1α CNX-TMD mutant displayed significantly more clusters than wild type IRE1α upon treatment with either Tg for 1 h and 2 h or Tm for 2 h (Figures 2G and 2H). However, both the wild type and IRE1α CNX-TMD mutant formed a similar number of clusters when cells were treated for a slightly longer period (4h) with Tm, but still the clusters were slightly bigger in cells expressing the mutant (Figures 2G and 2H). This result suggests that higher levels of ER stress can overcome Sec63-mediated inhibition of IRE1α clustering, which is consistent with our earlier studies (Sundaram et al., 2017). Taken together, these data suggest that IRE1α forms robust clusters in cells either deficient of Sec63, expressing Sec63 interaction defective mutants, or experiencing higher levels of ER stress.

### Sec63 attenuates IRE1α RNase activity in cells during persistent ER stress

Since Sec63 inhibits IRE1α clustering or higher-order oligomerization during ER stress, we wanted to determine if Sec63 also limits the RNase activity of IRE1α during ER stress conditions. To test this, we monitored IRE1α phosphorylation and its RNase-mediated splicing of XBP1 mRNA in both wild type and Sec63-/- cells treated with Tg. A significant proportion of IRE1α was activated after one hour of ER stress as represented by phosphorylated IRE1α (Figure 3A). IRE1α was mostly inactivated or dephosphorylated within eight hours of ER stress in wild type cells. The phosphorylation status of IRE1α was comparable with the IRE1α-mediated splicing of XBP1 mRNA splicing (Figure 3A and 3B). The ER stress-dependent BiP upregulation was also correlated with the inactivation of IRE1α in wild type cells. A small proportion of IRE1α was constitutively phosphorylated in Sec63-/- cells even under homeostatic conditions (Figure 3A). Upon ER stress, IRE1α was fully phosphorylated in Sec63-/- cells, but it could not be dephosphorylated or inactivated during the later hours of ER stress (Figure 3A and 3C). The continuous phosphorylation of IRE1α was correlated with its ability to mediate XBP1 mRNA splicing during ER stress. Even though BiP was highly upregulated in Sec63-/- cells (Figure S2D), it was not able to inactivate IRE1α in the absence of Sec63 (Figure 3A and 3C). The depletion of Sec63 did not appreciably affect the PERK branch of the UPR, as the ER stress-induced phosphorylation of PERK in Sec63-/- cells was similar to that of wild type cells (Figure S2A, S2B). The ATF6α branch was activated within one hour of ER stress in wild type cells as measured by the loss signal due to ER stress-induced proteolytic cleavage of ATF6α. Consistent with our previous studies (Sundaram et al., 2018), the ATF6α signal came back after eight hours of the treatment in wild type cells (Figures S2A). Surprisingly, the activation of ATF6α was significantly inhibited in Sec63-/- cells as shown by mostly full-length ATF6α during ER stress conditions (Figure S2A and S2C). However, ATF6α could be efficiently activated in Sec63-/- cells upon treatment with the strong ER stress inducer DTT (Figure S2D). This result suggests that ATF6α proteins are functional in Sec63-/- cells, but they require a strong ER stress inducer for their activation. IRE1α attenuation defects observed in Sec63-/- cells were not specific to cells treated with Tg since we obtained a similar result when cells were treated with Tm for up to 24 hours (Figure S3A). In contrast to Sec63-/- cells, the ER stress-dependent activation of IRE1α was inhibited in Sec62-/- cells (Figure S3B). This result suggests that IRE1α attenuation defects observed in Sec63-/- are not likely caused by inefficient protein translocation into the ER lumen since both Sec62 and Sec63 are involved in protein translocation.

**Figure 3.**
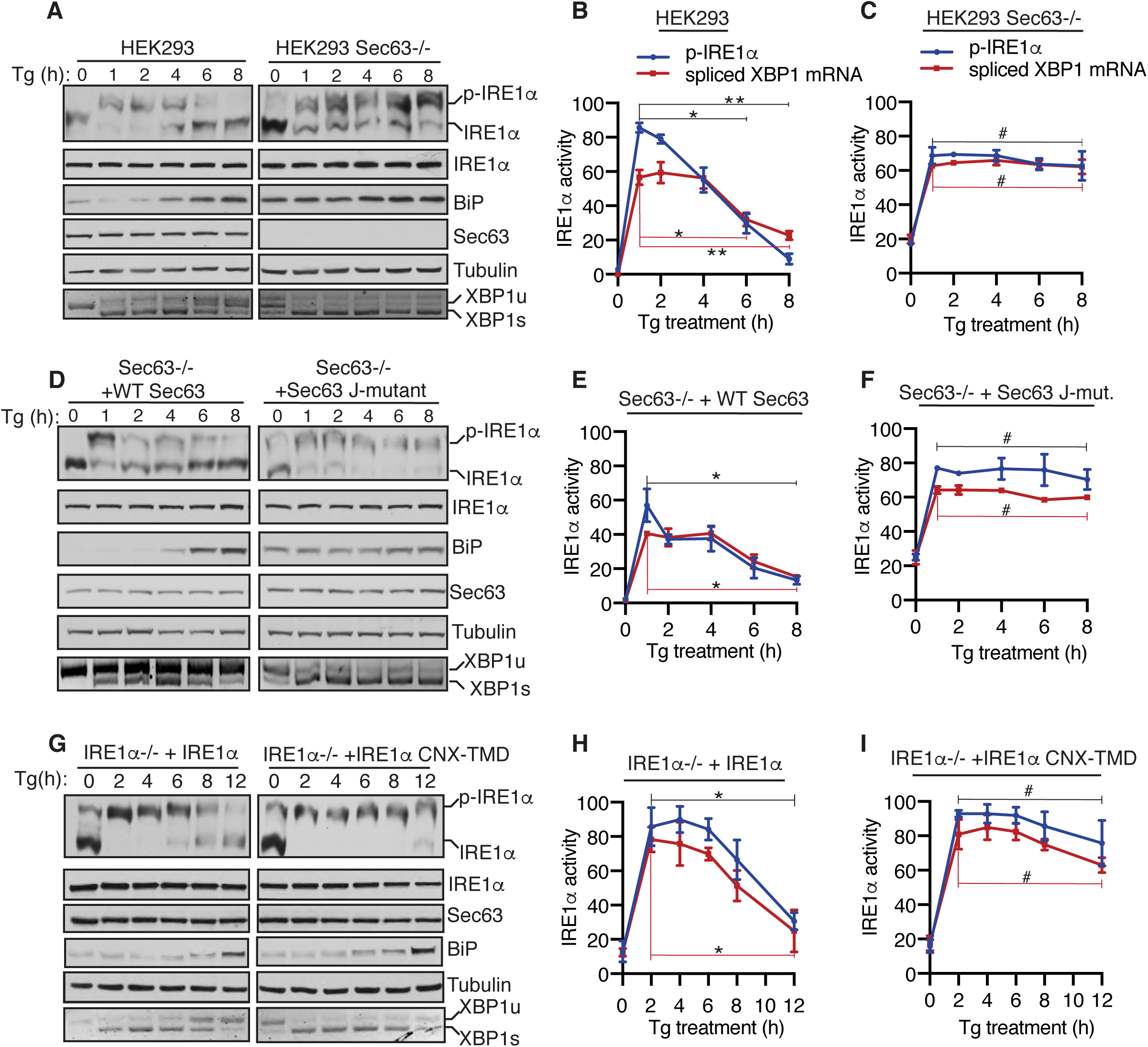
The J domain of Sec63 is essential for attenuating IRE1α activity during ER stress in cells. (A) Wild type HEK293 or Sec63-/- cells were treated with 2.5 μg/ml of Tg for the indicated time points and analyzed by immunoblotting as well as the XBP1 mRNA splicing assay. p-IRE1α denotes the phosphorylated form of IRE1α, which migrates slower in the phos-tag immunoblotting. XBP1u - Unspliced XBP1 mRNA, XBP1s - spliced XBP1 mRNA. (B, C) Quantification results of IRE1α phosphorylation and XBP1 mRNA splicing in (A). Error bars represent standard error of the mean (SEM) from three independent experiments. Statistical significance of IRE1α attenuation was compared between 1h time point (activated state) and the later time points (2, 4, 6, 8h) using Student’s t-test. Non-detectable (#) *P*>0.05, \**P*<0.05, \*\**P*<0.01. (D) Sec63-/- cells complemented with either WT or the J-domain mutant of Sec63 were treated and analyzed as in panel A. (E, F) Quantification results of IRE1α phosphorylation and XBP1 mRNA splicing in (D) were analyzed as in (B, C). (G) HEK293 IRE1α-/- cells stably complemented with either WT IRE1α or IRE1α-CNX-TMD were treated with 2.5 μg/ml of Tg for the indicated time points and analyzed as in panel A. (H, I) Quantification results of IRE1α phosphorylation and XBP1 mRNA splicing in (G) were analyzed as in (B, C). See also **Figure S2, S3, and S4**.

Next, we rescued IRE1α inactivation defects by complementing wild type Sec63 into Sec63-/- cells. The complementation of Sec63 partially restored activation and inactivation kinetics of IRE1α as mirrored by both IRE1α phosphorylation and XBP1 mRNA splicing (Figure 3D, 3E, and 3F). A proportion of IRE1α was constitutively activated even under homeostatic conditions in Sec63-/- cells complemented with the Sec63 J-domain mutant. Upon ER stress, IRE1α was efficiently activated in these cells but could not be attenuated during the later hours of ER stress, suggesting that the J-domain of Sec63 is required for inhibiting IRE1α activity during persistent ER stress. The activation and inactivation of PERK and ATF6α behaved quite similarly to Sec63-/- cells (Figure S4A, S4B, and S4C). We also complemented Sec63-/- cells with Sec63 mutants (Δ367-760 and Δ637-760) that have intact J-domains but weakly interacted with the Sec61 translocon (Figure S1B). These mutants failed to rescue IRE1α attenuation defects observed in Sec63-/- cells during ER stress (Figure S4D). This result implies that IRE1α must be close to Sec63 for efficient attenuation of its activity during ER stress.

Lastly, we tested the role of Sec63 in attenuating IRE1α activity using an approach that does not disrupt the function of Sec63 in cells. We therefore monitored IRE1α activity in cells expressing either wild type IRE1α or IRE1α CNX-TMD mutant, which weakly interacts with Sec61/Sec63 (Figure 1B). IRE1α CNX-TMD mutant was fully activated upon ER stress but displayed a significant defect in attenuation during persistent ER stress relative to wild type IRE1α as shown by both phosphorylation and XBP1 mRNA splicing (Figure 3G, 3H, and 3I). We noticed that a small fraction of IRE1α is activated in cells expressing either wild type IRE1α or IRE1α CNX-TMD mutant even under normal conditions (0h time point). This is presumably caused by a slight overexpression of complemented IRE1α in these cells (Sundaram et al., 2018). The activation and inactivation profiles of PERK and ATF6 in cells expressing IRE1α CNX-TMD mutant were identical to cells expressing wild type IRE1α (Figure S4E, S4F, and S4G). Taken together, our data establish that IRE1α inactivation was significantly impaired during persistent ER stress, either in the absence of Sec63 or when it failed to interact with Sec63.

### The Sec61/Sec63 complex recruits BiP to bind onto IRE1α in cells and *in vitro*

Next, we investigated the mechanism by which Sec63 turns off IRE1α activity during persistent ER stress. We hypothesized that Sec63 may recruit and activate the BiP ATPase via its luminal J-domain to bind onto IRE1α, leading to inhibition of IRE1α oligomerization and activity. To this end, we first tested whether the endogenous IRE1α/Sec61/Sec63 complex binds to BiP. The affinity purification of the chromosomal 2XStrep-tagged Sec61α from HEK293 cells revealed the endogenous complex of Sec61, Sec63, IRE1α, and BiP (Figure 4A). The reciprocal pull-down of the endogenous IRE1α using IRE1α antibodies further demonstrated the existence of the endogenous IRE1α/Sec61/Sec63/BiP complex in cells (Figure 4B).

**Figure 4.**
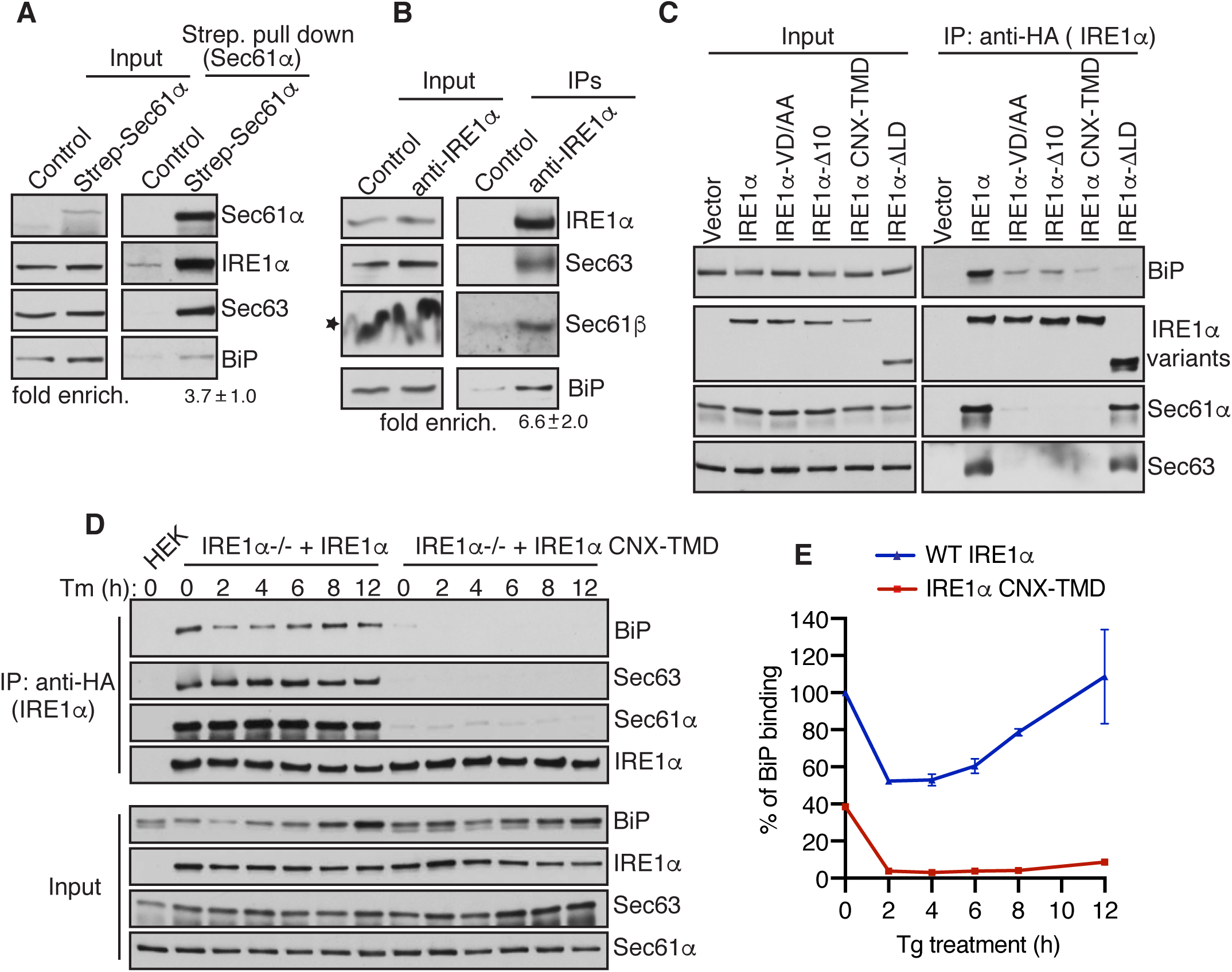
IRE1α binding to BiP strictly depends on its interaction with Sec61/Sec63 in cells. (A) Control HEK293 cells or chromosomal 2x-Strep-tagged Sec61α HEK293 cells were lysed and affinity purified using Strep-Tactin beads. The bound proteins were eluted and analyzed by immunoblotting for the indicated antigens. Fold enrichment of BiP (\*\**p* < 0.01) over the control was provided underneath the immunoblot. (B) HEK293 cells were lysed and immunoprecipitated using either control anti-His antibodies or anti-IRE1α antibodies. The immunoprecipitates were analyzed by immunoblotting for the indicated antigens. Fold enrichment of BiP (\**p*< 0.05) over the control was provided underneath the immunoblot. Star indicates that the distortion of the Sec61β band in the input was caused by co-migration with a high concentration of digitonin in the lysis buffer. (C) HEK293 cells were transiently transfected with either IRE1α-HA or its variants and immunoprecipitated using an anti-HA antibody and analyzed by immunoblotting. (D) HEK293 IRE1α-/- cells stably expressing IRE1α-HA or IRE1α-CNX-TMD-HA were induced with 5ng/ml of doxycycline and left either untreated or treated with 10 μg/ml Tm for the indicated time points. The treated cells were harvested, lysed, and subjected to immunoprecipitation using anti-HA magnetic beads. The immunoprecipitates were analyzed by immunoblotting for the indicated antigens. (E) Quantification results from IRE1α binding to BiP in (D). Error bars represent standard error of the mean from two independent experiments. Wild type IRE1α binding to BiP in unstressed cells was set as 100%. See also **Figure S5**.

We then examined whether IRE1α binding to BiP depends on its interaction with the Sec61/Sec63 complex. Since Sec63-/- cells contain high levels of BiP relative to wild type cells (Figure S2D), it was difficult to compare BiP binding to IRE1α between these two different cell lines. We therefore took advantage of our various Sec61/Sec63 interaction defective IRE1α mutants (Figure 1B) and performed co-immunoprecipitation studies to monitor their interaction with BiP. Wild type IRE1α associated with BiP along with the Sec61/Sec63 complex, whereas the translocon interaction defective IRE1α mutants markedly reduced interaction with BiP (Figure 4C). IRE1α mutant that is deleted of the luminal domain (LD) showed very little binding to BiP, but its interaction with Sec61/Sec63 was mostly unaffected (Figure 4C). This result suggests that BiP specifically binds to IRE1α in this complex. This conclusion was further supported by the finding that IRE1α still bound to BiP when immunoprecipitations were performed using the NP40/Deoxycholate detergent lysis buffer, which disrupts the interaction between IRE1α and Sec61/Sec63 during the cell lysis (Figure S5A). The recruitment of BiP to IRE1α was also dependent on the J-domain of Sec63 since IRE1α binding to BiP was reduced in cells overexpressing the Sec63 J-domain mutant in cells compared to wild type Sec63 expressing cells (Figure S5B).

We next tested whether Sec63-mediated BiP binding to IRE1α is sensitive to ER stress. Immunoprecipitation of IRE1α revealed that BiP was dissociated from IRE1α under all ER stress conditions relative to non-treated cells (Figure S5C). As expected, BiP binding to the Sec61/Sec63 interaction defective IRE1α CNX-TMD mutant was significantly reduced even under unstressed conditions, and that the interaction was almost abolished upon treatment with ER stress inducers (Figure S5C). Since Sec63 plays a crucial role in attenuation IRE1α activity during persistent ER stress conditions (Figure 3A, 3B), we hypothesized that Sec63 may also mediate complex formation between IRE1α and BiP during persistent ER stress, which would result in de-oligomerization and attenuation of IRE1α RNase activity. In support of this hypothesis, IRE1α binding to BiP was reduced upon acute ER stress, but the interaction was restored during persistent ER stress (Figure 4D and 4E). By contrast, IRE1α CNX-TMD mutant was not able to bind BiP under all conditions, supporting the conclusion that IRE1α binding to BiP strictly depends on its interaction with Sec63.

Next, we asked whether Sec61/Sec63 is sufficient to mediate BiP binding to IRE1α. To address this, we purified the IRE1α/Sec61/Sec63 complex from HEK293 cells stably expressing 2x Strep-tagged IRE1α-FLAG. A coomassie stained gel revealed that IRE1α was about three times more than Sec61/Sec63 since the complex was purified from cells overexpressing IRE1α (Figure 5A). As a control, we similarly purified IRE1α Δ10, which lacks the interaction with the Sec61/Sec63 complex. We expressed and purified recombinant BiP from E. coli (Figure 5B). We first prepared anti-FLAG antibody beads bound to the IRE1α complex or IRE1α Δ10. We then incubated the beads with or without BiP in the presence or absence of ATP. In the absence of ATP, BiP bound to both the IRE1α/Sec61/Sec63 complex and IRE1α Δ10 (Figure 5C). BiP was mostly dissociated from IRE1α Δ10 in the presence of ATP (Figure 5C), likely due to ATP bound BiP has higher substrate dissociation rates (Misselwitz et al., 1998). In sharp contrast, BiP binding to IRE1α/Sec61/Sec63 was intact even in the presence of ATP (Figure 5C). This result suggests that the J-domain of Sec63 stimulated ATP hydrolysis of BiP to bind onto IRE1α. We also obtained a similar result of Sec61/Sec63 dependent BiP binding onto IRE1α when the components were incubated in solution followed by immunoprecipitation with anti-FLAG beads (Figure S6). Taken together our in vitro results suggest that Sec61/Sec63 is sufficient to mediate BiP binding to IRE1α in the presence of ATP.

**Figure 5.**
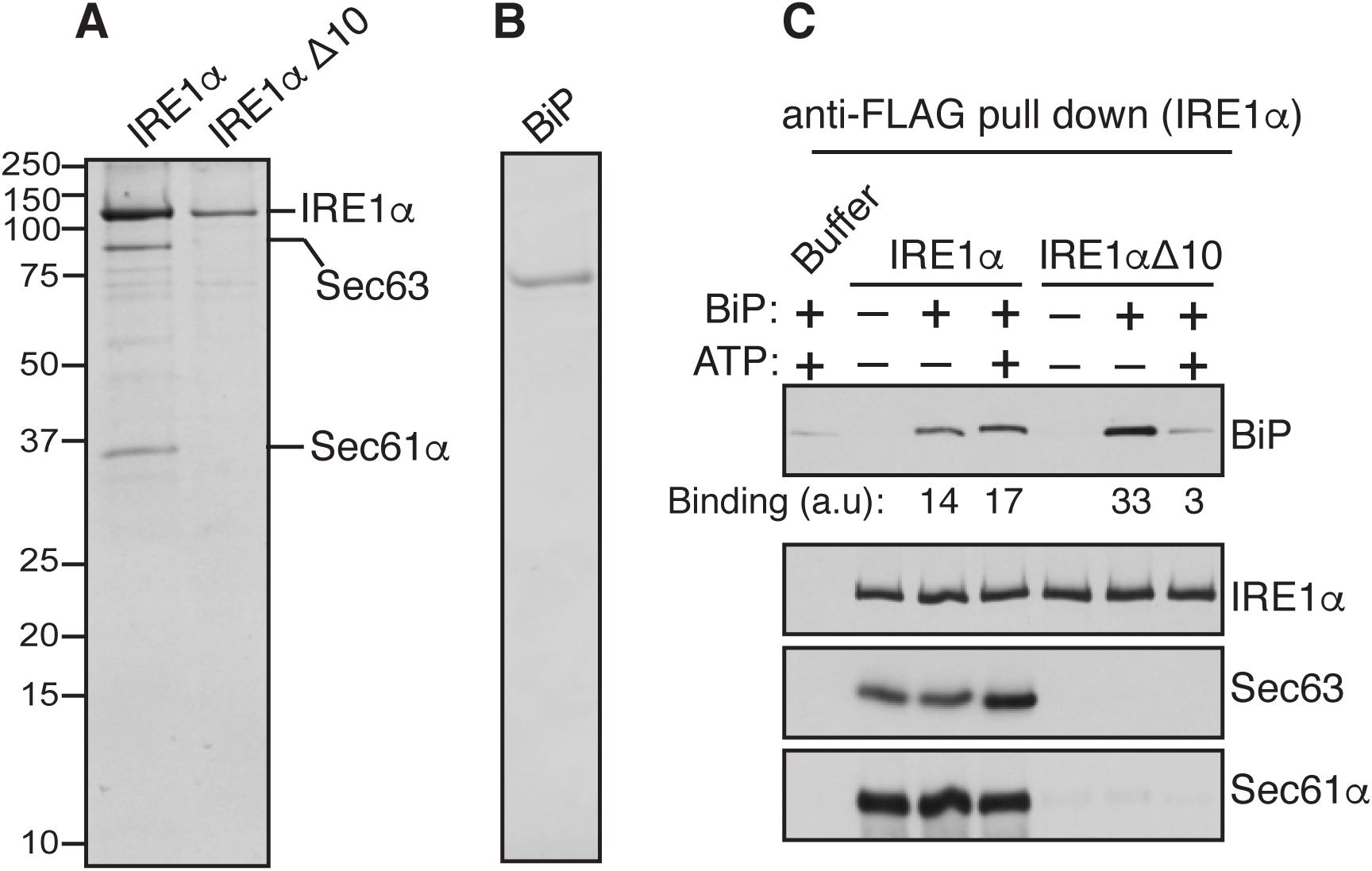
Biochemical reconstitution of Sec61/Sec63-mediated BiP binding to IRE1α. (A) A coomassie blue stained gel showing the purified IRE1α/Sec61/Sec63 complex or IRE1α Δ10 from HEK293 stably expressing 2xStrep-IRE1α-FLAG or 2xStrep-IRE1α Δ10-FLAG. (B) A coomassie blue stained gel showing purified His-BiP from E. coli. (C) The purified IRE1α/Sec61/Sec63 complex or IRE1αΔ10 was bound to anti-FLAG beads and incubated with or without BiP in the presence or absence of ATP as shown. After incubation, IRE1α bound anti-FLAG beads were washed and eluted with sample buffer. A negative control reaction was performed by incubating empty anti-FLAG beads with the buffer, BiP, and ATP. The samples were analyzed by immunoblotting for the indicated antigens. BiP bands were quantified and presented as arbitrary units (a.u) after subtracting the buffer background. See also **Figure S6**.

## Discussion

It is known that IRE1α RNase activity is attenuated during persistent ER stress (Lin et al., 2007). The failure to attenuate IRE1α activity during prolonged ER stress can induce cell death, which is linked to many human diseases (Ghosh et al., 2014; Hetz, 2012; Lin et al., 2007). In this study, we identify the translocon associated membrane protein Sec63 as a critical factor that turns off of IRE1α activity during persistent ER stress. We show that Sec63 recruits and activates BiP ATPase via its luminal J-domain to bind onto IRE1α, thus suppressing higher-order oligomerization and RNase activity of IRE1α during ER stress (Figure 6). We envision that defects in attenuation of IRE1α signaling during ER stress may be detrimental to cells burdened with high levels of secretory proteins such as pancreatic beta cells (Back and Kaufman, 2012).

**Figure 6.**
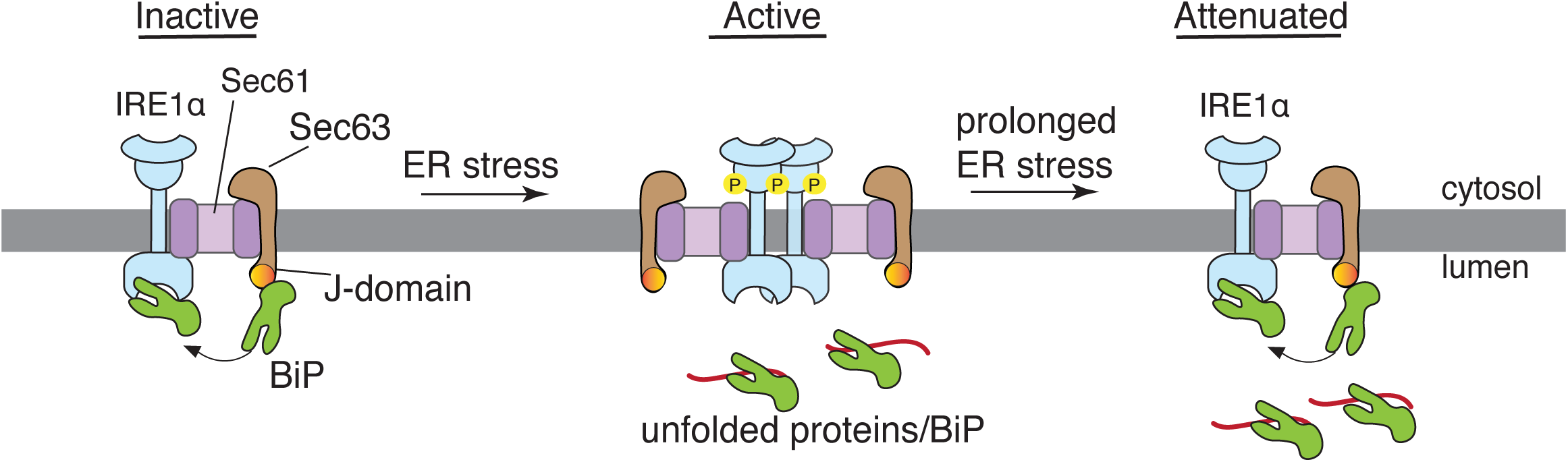
Model of the Sec61/Sec63/BiP-mediated attenuation of IRE1α signaling during prolonged ER stress. The J-domain of Sec63 recruits and activates BiP to bind onto IRE1α under normal conditions. Upon ER stress, BiP is released from IRE1α to bind to unfolded proteins, leading to IRE1α oligomerization and activation. Once activated IRE1α-mediates XBP1 mRNA splicing, resulting in the production of the active transcription factor XBP1s that activates the expression of ER chaperones and quality control factors (not shown). During prolonged ER stress, Sec63 recruits the upregulated BiP to bind onto IRE1α and hence inhibiting IRE1α oligomerization and RNase activity.

It has long been known that BiP plays a central role in regulating the oligomerization and activity of IRE1α, (Amin-Wetzel et al., 2017; Carrara et al., 2015; Oikawa et al., 2009; Preissler and Ron, 2019; Ricci et al., 2019). However, it is unclear how the luminal BiP is recruited to the membrane-localized IRE1α, which is extremely low abundant (Kulak et al., 2014), to inhibit IRE1α oligomerization even during ER stress. Our previous studies have shown that most of the endogenous IRE1α proteins exist in a complex with the Sec61 translocon complex (Plumb et al., 2015). In this study, our data establish that Sec63 is a part of the IRE1α/Sec61 translocon complex. Our interaction studies suggest that the Sec61 translocon bridges the interaction between IRE1α and Sec63. Future structural studies are required to determine the precise arrangement of this complex. Although Sec62 is known to associate with Sec61/Sec63, it is not enriched in IRE1α immunoprecipitates, suggesting that IRE1α selectively interacts with a Sec61 translocon complex that contains Sec63 but not sec62. Another possibility is that Sec62 interaction with the IRE1/Sec61/Sec63 complex is labile during immunoprecipitations. This is consistent with the previous studies that Sec62 interaction with the Sec61 translocon complex is sensitive to high salt concentrations (Meyer et al., 2000).

Depletion of Sec63 but not Sec62 induces the formation of IRE1α higher-order oligomers or clusters in cells upon ER stress. Specifically, the J domain of Sec63 is required for suppressing IRE1α clusters. This is further supported by the observation that the Sec61/Sec63 interaction defective mutants readily form clusters in cells upon ER stress (Sundaram et al., 2017). Our data suggest that increased levels of IRE1α clusters in Sec63 deficient cells correlate with attenuation defects of IRE1α observed in these cells. By contrast, Sec62 is dispensable for regulating IRE1α oligomerization and RNase activity because it neither contains J-domain nor is required for the stability of Sec63 in cells. Surprisingly, the ER stress-dependent activation of IRE1α was significantly inhibited in Sec62-/- cells. One explanation for this observation is that since Sec63 remains intact in these cells, it can efficiently recruit BiP, which is highly upregulated in Sec62-/- cells, and suppress the activation of IRE1α even during ER stress.

Two lines of evidence support that Sec63 is responsible for recruiting luminal BiP to bind onto IRE1α. First, IRE1α mutants that weakly interact with Sec61/Sec63 show a significantly less binding to BiP. It is unlikely that these IRE1α mutants also disrupt their interaction with other luminal J-domain containing proteins because the Sec61/Sec63 interacting regions are localized in both luminal juxtamembrane and transmembrane regions of IRE1α, which should not interfere with IRE1α interaction with soluble luminal proteins. Second biochemical reconstitution experiments with purified proteins suggest that Sec61/Sec63 is sufficient to mediate BiP binding to IRE1α in the presence of ATP. Although BiP binding to IRE1α/Sec61/Sec63 is persistent in the presence of ATP, the binding is not significantly increased compared to the condition without ATP. This is likely due to the sub-stoichiometric amount of Sec63 (∼3X less) compared to IRE1α in our in vitro reactions. By contrast, the concentration of Sec63 is vastly more abundant than IRE1α in cells (Kulak et al., 2014). Also, the presence of detergent, which is added to keep the membrane proteins soluble, in reactions may disrupt the efficient binding of BiP to IRE1α. Future studies will focus on reconstituting the IRE1α/Sec61/Sec63 complex into a supported bilayer and monitor the BiP-mediated de-oligomerization of IRE1α.

The ER contains seven J-domain containing proteins localized in the ER lumen where they can interact with BiP (Pobre et al., 2019). It is conceivable that other J-domain containing proteins may be involved in the attenuation of IRE1α activity. A recent study suggests that ERdj4 is required for complex formation between IRE1α and BiP, thus repressing IRE1α oligomerization and activation (Amin-Wetzel et al., 2017). However, our data suggest that ERdj4 or other J-domain containing proteins may not play a significant role in attenuation of IRE1α because the majority of IRE1α remains activated in Sec63 knockout cells during persistent ER stress. It is unlikely that the depletion of Sec63 affects the translocation of ERdj4 and thereby circumventing the alternative pathway for IRE1α attenuation. This is because IRE1α CNX-TMD mutant that weakly interacts with Sec61/Sec63 also show a significant defect in their attenuation during persistent ER stress. Although IRE1α CNX-TMD interaction with BiP was almost abolished, it is not significantly activated in unstressed conditions but fully activated upon ER stress. This result suggests that the release of BiP from IRE1α alone is not sufficient for efficient activation of IRE1α in unstressed conditions. Instead, the presence of misfolded proteins induced by ER stress is critical for efficient activation of IRE1α in cells.

Since the Sec61 translocon selectively associates with the IRE1α branch of the UPR, depletion of Sec63 has little effects on activation PERK. This is consistent with previous studies that either depletion of Sec63 or Sec61 activated IRE1α but not PERK (Adamson et al., 2016; Fedeles et al., 2015). Our data show that ATF6α activation is significantly inhibited in Sec63-/- cells upon ER stress. We speculate that this is caused by the presence of excess BiP in these cells, which may effectively suppress ATF6α activation. This is supported by our previous studies that the overexpression of recombinant BiP in cells mostly inhibits the activation of ATF6α but has little effects on the activation of PERK during ER stress (Sundaram et al., 2018). Since IRE1α but not PERK and ATF6α associates with the translocon complex, it may also function as a sensor for protein translocation into the ER lumen. We speculate that IRE1α may be activated when the translocating substrate either sequesters BiP from IRE1α and/or directly binds to the luminal domain of IRE1α. This preemptive mechanism would optimize the protein folding capacity of the ER based on individual substrates that depend on BiP for their maturation.

Recent structural studies suggest that Sec63 binding to the translocon sterically hinders the ribosome binding to the translocon (Itskanov and Park, 2019; Wu et al., 2019). Future studies are warranted to determine whether Sec63 is still associated with the translocon when the ribosome-nascent chain complex is recruited to the Sec61/IRE1α complex. Intriguingly, a recent study shows that IRE1α can directly bind to ribosomes (Acosta-Alvear et al., 2018), suggesting that IRE1α forms an intricate complex with the translocon-ribosome complex.

Structural and biochemical studies are needed to visualize this complex to understand how IRE1α monitors and controls protein translocation into the ER.

## STAR*METHODS

Detailed methods are provided in the online version of this paper and include the following:

## KEY RESOURCES TABLE

### RESOURCE AVAILABILITY

Lead Contact

Material Availability

Data and Code Availability

## EXPERIMENTAL MODEL AND SUBJECT DETAILS

Cell culture

## METHOD DETAILS

DNA constructs

CRISPR/Cas9-mediated knockout or knock-in cell lines

Generation of stable cell lines

Immunoprecipitations

Isolation of the endogenous IRE1/Sec61/Sec61/BiP complex

Purification of the IRE1α complex and BiP

In vitro reconstitution Sec61/Sec63-mediated BiP binding to IRE1α

Immunofluorescence

XBP1 mRNA splicing assay

Phostag-based immunoblotting

Quantification and statistical analysis

## SUPPLEMENTAL INFORMATION

Supplemental Information can be found online at https:

## ACKNOWLEDGEMENTS

We thank Dr. Ramanujan Hegde for Sec62 and Sec63 antibodies, and Dr. Stefan Somlo for mouse Sec63 plasmid. We thank Jacob Culver for useful discussion and comments on the manuscript. This work is funded by NIH grants R01GM117386 (M.M) and R21AG056800 (M.M).

## AUTHOR CONTRIBUTIONS

XL designed and performed most of the experiments. SS generated Sec63 constructs and purified BiP protein. MM performed BiP binding to IRE1α in vitro, SA created Sec63-/- and 2XStrep-tagged Sec61α HEK293 cells as well as performed initial experiments, RP characterized the IRE1α CNX-TMD mutant and established cell lines. AS purified the IRE1α/Sec61/Sec3 complex. MM supervised the project and wrote the manuscript with inputs from all the authors.

---

**Figure S1, related to Figure 1.**
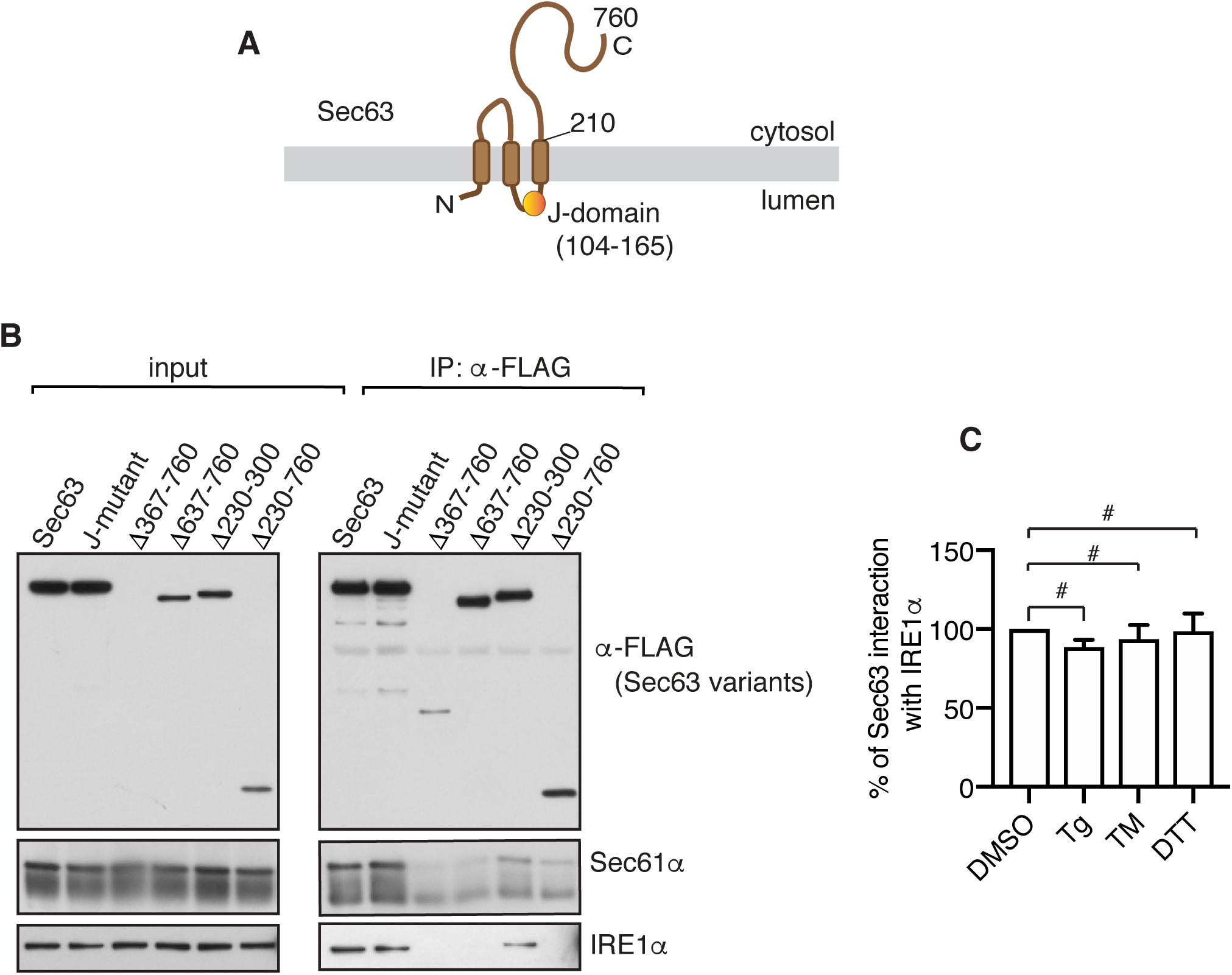
IRE1α interacts with Sec63 through the Sec61 translocon. (A) A diagram showing the topology of Sec63. (B) The cell lysates of the indicated versions of FLAG-tagged Sec63 were immunoprecipitated with anti-FLAG beads and analyzed by immunoblotting for the indicated antigens. The HPD tripeptide in the J-domain was replaced with AAA to create the J-domain mutant of Sec63. (C) Quantification results of Figure 1E. Error bars represent the standard error of the mean (SEM) from three independent experiments. p-value higher than 0.05 is represented by # (non-detectable). The percentage of IRE1α bound to Sec63 in the DMSO treated sample was set as 100%.

**Figure S2, related to Figure 3.**
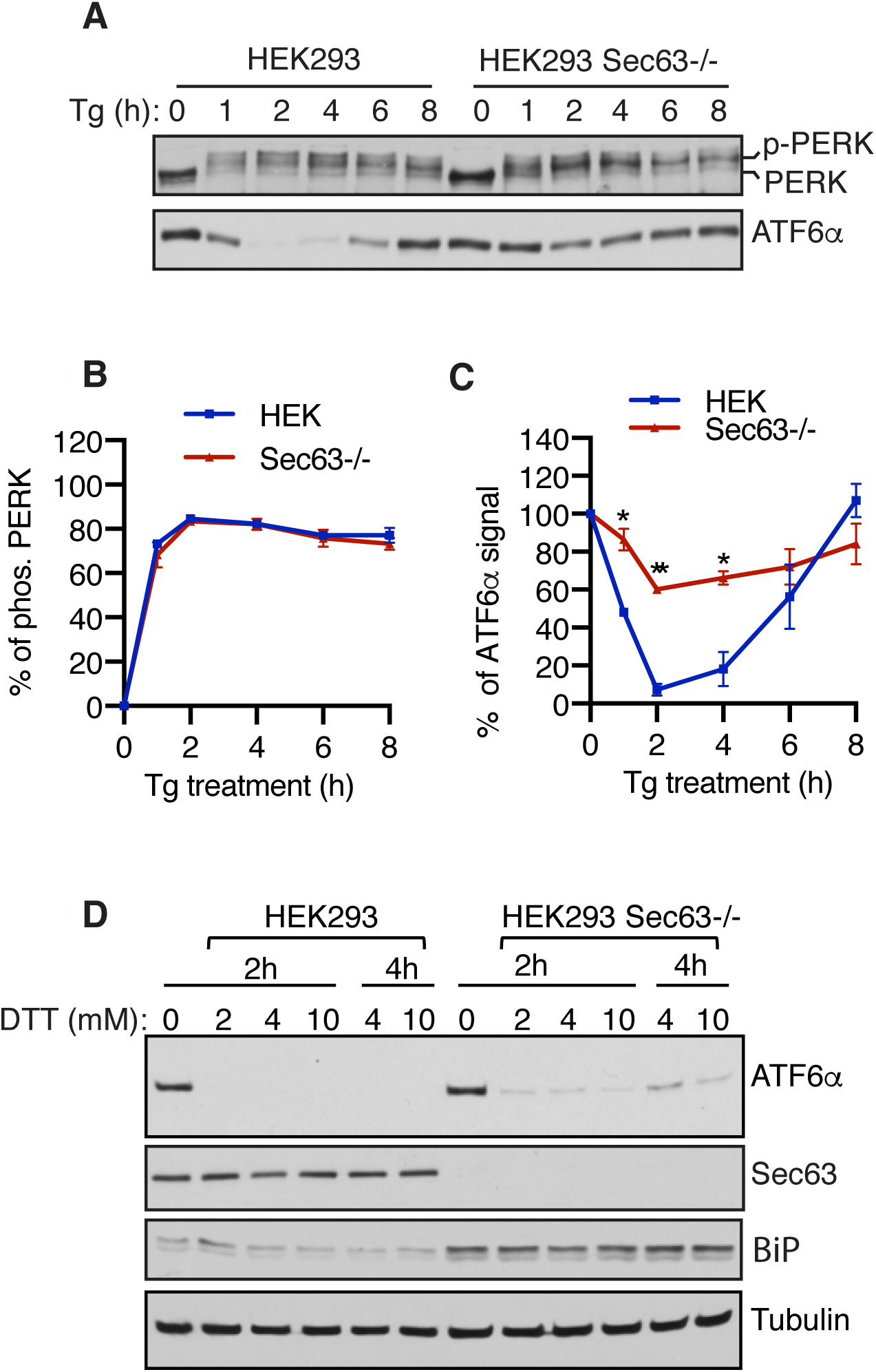
Probing activation and inactivation of PERK and ATF6α in Sec63-/- cells. (A) Wild type HEK293 or Sec63-/- cells were treated with 2.5 μg/ml of Tg for the indicated time points and analyzed by immunoblotting for PERK and ATF6α. The ATF6α bands represent “full length” proteins. The loss of ATF6α signals indicates the cleavage of its N-terminal cytosolic domain during ER stress. Our ATF6α antibodies were not suitable for detecting the cleaved N-terminal cytosolic domain. (B) Quantification results of PERK phosphorylation in (A). Error bars represent the standard error of the mean (SEM) of three independent experiments. PERK phosphorylation difference in WT cells and Sec63-/- cells was determined by Student’s t test. p values were non-significant, as they are higher than 0.05. (C) Quantification results of ATF6α signal in (A). Error bars represent the standard error of the mean (SEM) of three independent experiments. ATF6α signal difference in WT cells and Sec63-/- cells was determined by Student’s t-test. *P<0.05; **P<0.01. (D) HEK293 or Sec63-/- cells were treated with DTT for the indicated time points with the indicated concentrations. The cells were directly harvested in SDS sample buffer and analyzed by immunoblotting for the indicated antigens.

**Figure S3, related to Figure 3.**
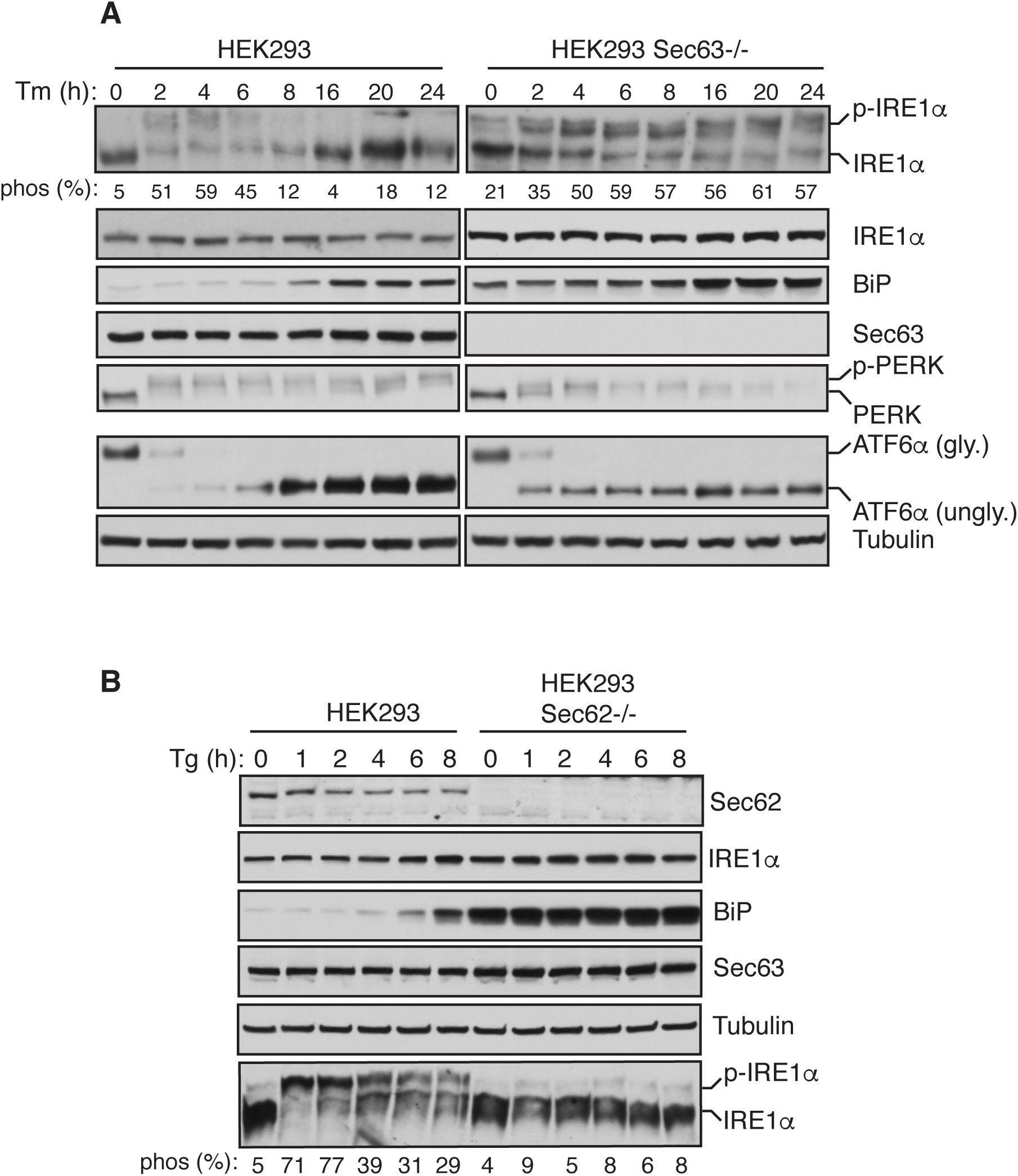
IRE1α activation and inactivation in Sec63-/- or Sec62-/- cells. (A) Wild type HEK293 or Sec63-/- cells were treated with 5 μg/ml of Tm for the indicated time points and analyzed by immunoblotting for the indicated antigens. The percentage of IRE1α phosphorylation is shown underneath phos-tag immunoblots. ATF6 is glycosylated (gly.), but it is unglycosylated (ungly.) during the tunicamycin treatment. (B) Wild type HEK293 or Sec62-/- cells were treated with 2.5 μg/ml of Tg for the indicated time points and analyzed as in panel A.

**Figure S4, related to Figure 3.**
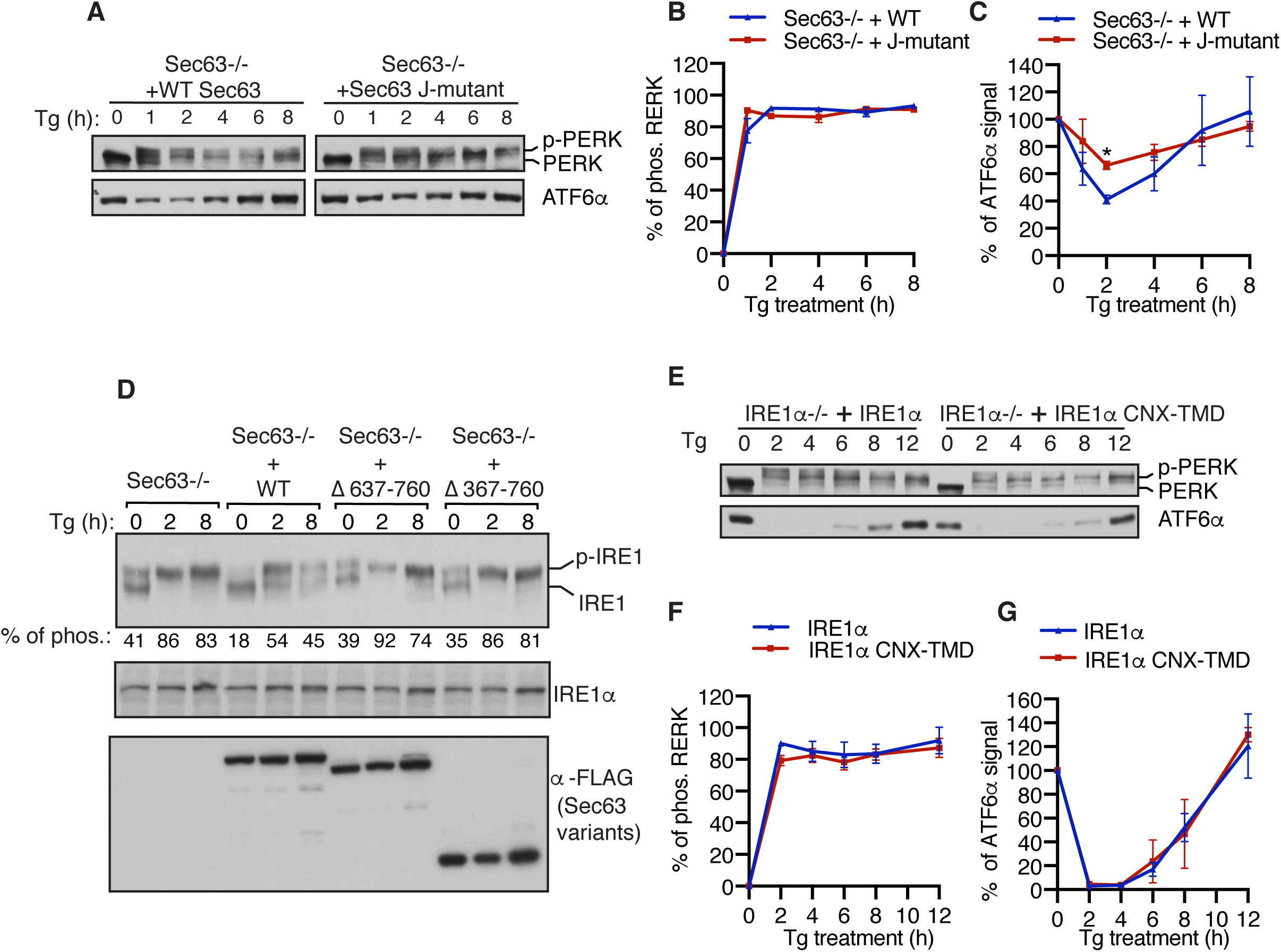
Probing activation and inactivation of IRE1α, PERK, and ATF6α in complemented cell lines. (A) Sec63-/- cells complemented with either WT or the J-domain mutant were treated with 2.5 μg/ml of Tg for the indicated time points and analyzed by immunoblotting for PERK and ATF6α. (B) Quantification results of PERK phosphorylation in (A). Error bars represent the standard error of the mean (SEM) of three independent experiments. PERK phosphorylation difference in WT cells and Sec63-/- cells was determined by Student’s *t* test. p values were non-significant, as they are higher than 0.05. (C) Quantification results of ATF6α signal in (A). Error bars represent the standard error of the mean (SEM) of three independent experiments. ATF6α cleavage difference in WT cells and Sec63-/- cells was determined by Student’s *t* test. *P<0.05. (D) Sec63-/- cells transiently expressing either WT or Sec63 mutants were treated with 2.5 μg/ml of Tg for the indicated time points and analyzed by immunoblotting. The percentage of IRE1α phosphorylation is shown underneath phos-tag immunoblots (E) IRE1α -/- cells complemented with either WT or IRE1α CNX-TMD were treated with 2.5 μg/ml of Tg for the indicated time points and analyzed by immunoblotting for PERK and ATF6α. (F) PERK phosphorylation in (E) was quantified as in (B). (G) ATF6α signal in (E) was quantified as in (C).

**Figure S5, related to Figure 4.**
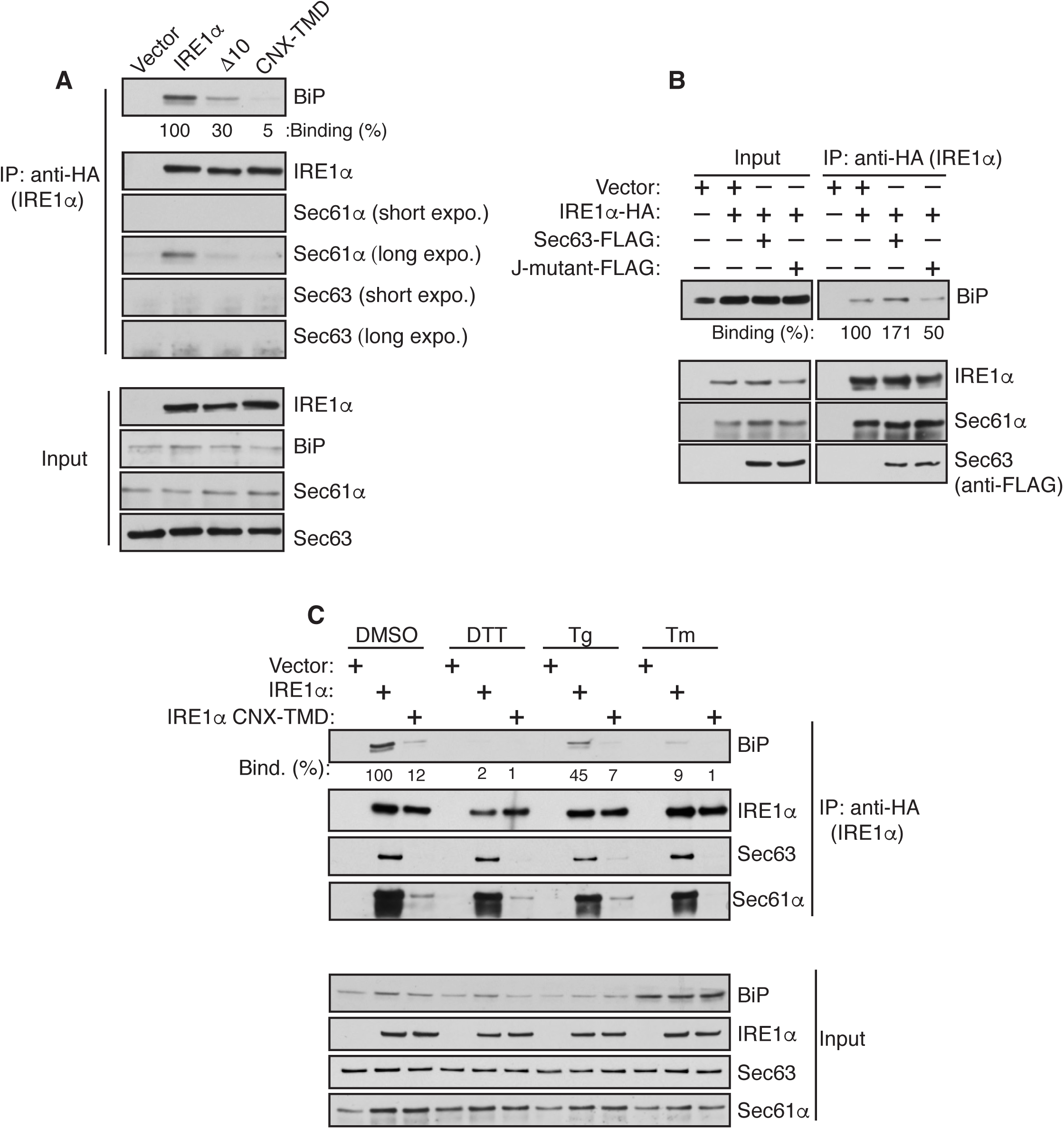
Sec61/Sec63-mediated BiP binding to IRE1α in cells. (A) HEK293 Sec63-/- cells complemented with wild type Sec63 were transiently transfected with either IRE1α-HA or its variants. The cell lysates were prepared using the buffer containing NP40/deoxycholate followed by immunoprecipitation with anti-HA magnetic beads and analyzed by immunoblotting for the indicated antigens. The amount of BiP binding to wild type IRE1α was taken as 100%. (B) HEK293 cells were co-transfected with IRE1α-HA and empty vector or Sec63-FLAG or J-domain mutant of Sec63-FLAG. The cell lysates were immunoprecipitated with an anti-HA antibody and analyzed by immunoblotting for the indicated antigens. The amount of BiP binding to IRE1α that was co-transfected with empty vector was taken as 100%. (C) HEK293 IRE1α-/- cells stably expressing IRE1α-HA or IRE1α-CNX-TMD-HA were treated with DMSO for 2 h, 4 mM DTT for 2 h, 5 μg/ml Tg for 2 h, or 10 μg/ml Tm for 4 h. The treated cells were harvested and analyzed as in A. The amount of BiP binding to wild type IRE1α in unstressed cells was taken as 100%.

**Figure S6, related to Figure 5.**
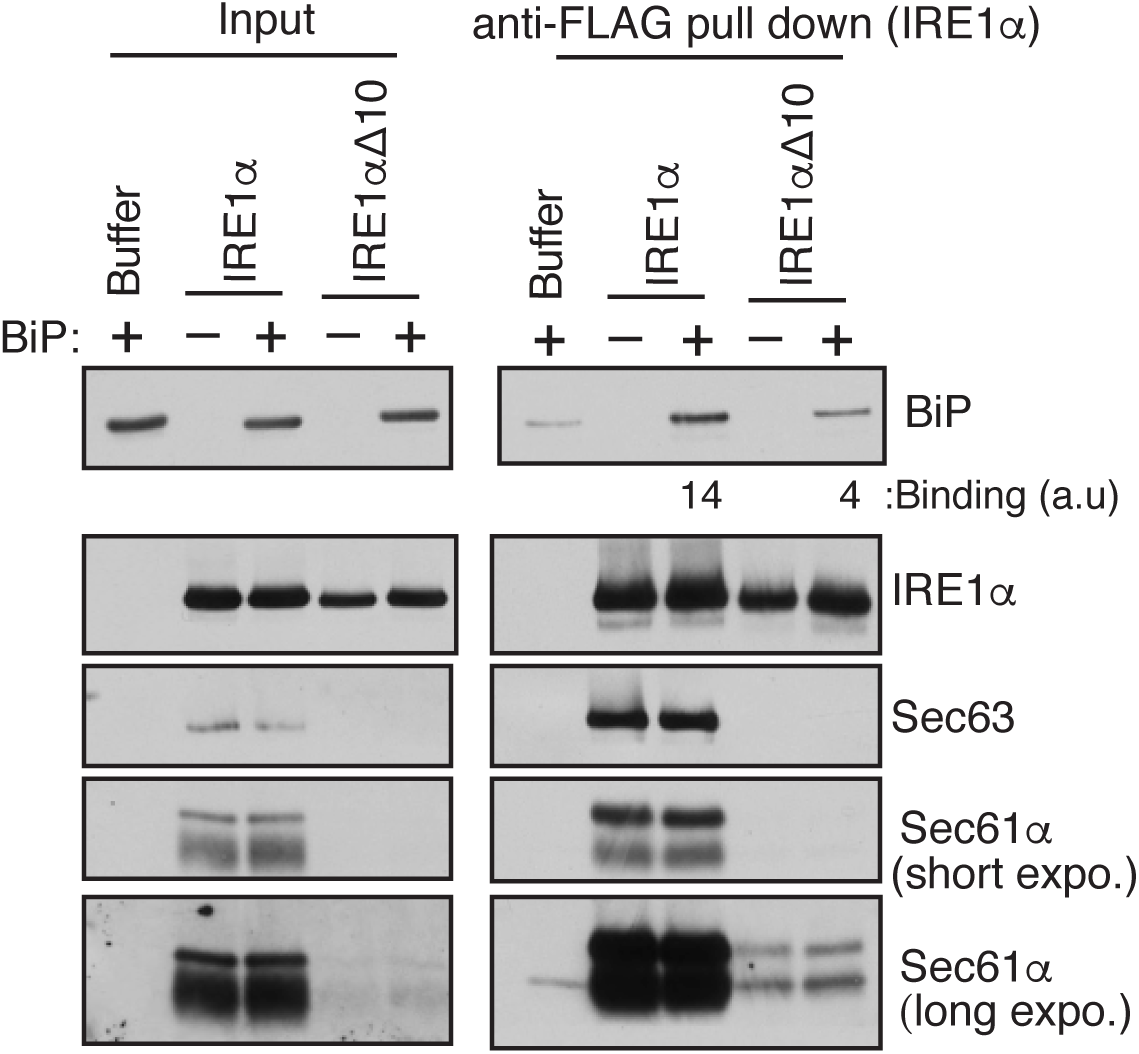
Sec61/Sec63-mediated BiP binding to IRE1α *in vitro*. The purified IRE1α/Sec61/Sec63 complex or IRE1αΔ10 was incubated with or without BiP in the presence of ATP. After incubation, IRE1α was immunoprecipitated using anti-FLAG beads. A negative control reaction was performed by mixing the buffer, BiP, and ATP, followed by immunoprecipitation with anti-FLAG beads. The samples were analyzed by immunoblotting for the indicated antigens. Note that IRE1α Δ10 contains a residual amount of the Sec61 translocon (long exposure blot), suggesting that weak binding between BiP and IRE1α Δ10 may be due to the presence of a small amount of Sec61/Sec63 in the sample. BiP bands were quantified and presented as arbitrary units (a.u) after subtracting the buffer background.

## STAR★Methods

KEY RESOURCES TABLE

| REAGENT or RESOURCE | SOURCE | IDENTIFIER |
| --- | --- | --- |
| **Antibodies** |  |  |
| Rabbit anti-IRE1α | Cell Signaling | 3294 |
| Rabbit anti-IRE1α | Santa Cruz | sc-20790 |
| Mouse anti-Tubulin | Abcam | ab7291 |
| Rabbit anti-BiP | Cell Signaling | 3177 |
| Mouse anti-BiP | BD Biosciences | 610979 |
| Rabbit anti-ATF6α | Cell Signaling | 65880 |
| Rabbit anti-PERK | Cell Signaling | 3192 |
| Goat anti-mouse IgG-HRP | Jackson ImmunoResearch | 11-035-003 |
| Goat anti-rabbit IgG-HRP | Jackson ImmunoResearch | 111-035-003 |
| Goat anti-rabbit IgG-Cy3 | Jackson Immuno Research | 711-165-152 |
| Rabbit anti-Sec61α | Gift from Dr. Ramanujan Hegde | N/A |
| Rabbit anti-Sec62 | Gift from Dr. Ramanujan Hegde | N/A |
| Rabbit anti-Sec63 | Gift from Dr. Ramanujan Hegde | N/A |
| Rabbit anti-HA | Gift from Dr. Ramanujan Hegde | N/A |
| Rabbit anti-Sec61β | Gift from Dr. Ramanujan Hegde | N/A |
| **Resins** |  |  |
| mouse anti-HA magnetic beads | Fisher Scientific | 88836 |
| Rat anti-FLAG beads | Biolegend | 651503 |
| StrepTactin-XT Sepharose | IBA, Germany | 2-1201-010 |
| Protein A agarose | Repligen | CA-PRI-1000 |
| **Reagents** |  |  |
| DMEM | Corning | 10-013-CV |
| FBS | Gibco | 16000044 |
| Horse Serum | Sigma | H0146 |
| Penicillin/Streptomycin | Gibco | 15140122 |
| Lipofectamine 2000 | Invitrogen | 11668019 |
| Doxycycline | Clontech | 631311 |
| Hygromycin B | Invitrogen | 10687010 |
| Blasticidin | InvivoGen | ant-bl-1 |
| Thapsigargin | Enzo Life Sciences | BML-PE180 |
| Protease inhibitor cocktail | Roche | 11873580001 |
| Poly L-lysine | Peptides International | OKK-3056 |
| Digitonin | EMD Millipore | 300410 |
| Fluoromount G | SouthernBiotech | 0100-01 |
| Phos-tag | Wako | 300-93523 |
| West Pico Substrate | Thermo Scientific | 34080 |
| Femto Substrate | Thermo Scientific | 34095 |
| Oligo(dT) primers | Qiagen | 79237 |
| M-MLV reverse transcripts | NEB | M0253 |
| Trizol reagent | Thermo Scientific | 15596026 |
| DMSO | Sigma-Aldrich | D2650 |
| Tunicamycin | Sigma- Aldrich | T7765 |
| DTT | AmaricanBio | 3483-12-2 |
| PonceauS solution | Sigma-Aldrich | P3504 |
| Sec62 siRNA | Qiagen | 68875 |
| Sec63 siRNA | Qiagen | 68711 and 68715 |
| **Experimental Models: Cell line** |  |  |
| HEK 293-Flp-In T-Rex cells | Invitrogen | R78007 |
| HEK 293-Flp-In Sec63-/- | (Sun and Mariappan, 2020) | N/A |
| HEK 293-Flp-In Sec62-/- | This study | N/A |
| HEK 293-Flp-In Sec63-/-+Sec63-FLAG | (Sun and Mariappan, 2020) | N/A |
| HEK 293-Flp-In Sec63-/-+Sec63 J-mutant-FLAG | This study | N/A |
| HEK 293-Flp-In IRE1α-/-+IRE1α-HA | (Plumb et al., 2015) | N/A |
| HEK 293-Flp-In IRE1 α-/-+IRE1α-CNX-TMD-HA | This study | N/A |
| 2XStrep-Sec61α | This study | N/A |
| **Recombinant DNA** |  |  |
| pOG44 | Invitrogen | V600520 |
| pSpCas9(BB)-2A-Puro | Addgene | 62988 |
| pCR-BluntII | Addgene | K2800-20 |
| pCDNA5/FRT/TO-empty vector | Invitrogen | V652020 |
| pCDNA5/FRT/TO-Sec63-Flag | (Sun and Mariappan, 2020) | N/A |
| pCDNA5/FRT/TO-Sec63 J mut-Flag | This study | N/A |
| pCDNA5/FRT/TO-IRE1α-HA | (Plumb et al., 2015) | N/A |
| pCDNA5/FRT/TO-IRE1α-HA | (Plumb et al., 2015) | N/A |
| pCDNA5/FRT/TO-IRE1α-CNX-TMD-HA | This study | N/A |
| pCDNA5/FRT/TO-IRE1α-VD/AA -HA | (Plumb et al., 2015) | N/A |
| pCDNA5/FRT/TO-IRE1α-ΔLD -HA | (Plumb et al., 2015) | N/A |
| pCDNA5/FRT/TO-IRE1α-M/A -HA | (Plumb et al., 2015) | N/A |
| pCDNA5/FRT/TO-IRE1α-L/A -HA | (Plumb et al., 2015) | N/A |
| pCDNA5/FRT/TO- Sec63 Δ367-760-FLAG | This study | N/A |
| pCDNA5/FRT/TO- Sec63 Δ637-760-FLAG | This study | N/A |
| pCDNA5/FRT/TO- Sec63 Δ230-300-FLAG | This study | N/A |
| pCDNA5/FRT/TO- Sec63 Δ230-760-FLAG | This study | N/A |

## RESOURCE AVAILABILITY

### Lead Contact

Further information and requests for resources and reagents should be directed to and will be fulfilled by the Lead Contact, Malaiyalam Mariappan.

### Materials Availability

All unique/stable reagents generated in this study are available from the corresponding author with a completed Materials Transfer Agreement.

### Data and Code Availability

Original data will be made available upon request to the lead contact.

## EXPERIMENTAL MODEL AND SUBJECT DETAILS

HEK 293-Flp-In T-Rex cells and derivative cells were cultured in Dulbecco’s Modified Eagle’s Medium (DMEM) and 10% fetal bovine serum (FBS) and 100 U/mL penicillin and 100 µg/mL streptomycin at 5% CO2. IRE1α-/- or Sec63-/- HEK293 cells created by CRISPR/Cas9 were previously described (Plumb et al., 2015; Sun and Mariappan, 2020). Sec62-/- and 2X Strep-tagged Sec61α HEK293 cells were described in method details. Knockout and stable cell lines were routinely verified by immunoblotting.

## METHOD DETAILS

### DNA constructs

For mammalian cell expression, cDNAs were cloned into pcDNA5/FRT/TO (Invitrogen). Constructs encoding IRE1α-HA and its mutants were previously described (Plumb et al., 2015). The TMD of IRE1α was replaced with calnexin TMD to create IRE1α-CNX-TMD-HA using the protocol previously described (Volmer et al., 2013). Mouse Sec63 plasmid was a kind gift from Dr. Stefan Somlo (Yale School of Medicine). Sec63 truncation constructs, Δ367-760, Δ637-760, Δ230-300, and Δ230-760, were made using phosphorylated primers with the Phusion Site-Directed Mutagenesis protocol. The tripeptide HPD in the J-domain was replaced with AAA to create the J-domain mutant of Sec63 using site-directed mutagenesis. Rat BiP lacking the N-terminal signal sequence (1-18 amino acids) was cloned into pET-28a (+) using a standard cloning procedure. 3% DMSO was included in all PCR reactions to enhance amplification. The coding regions of all constructs were verified by sequencing performed in the Yale Keck DNA Sequencing Facility.

### CRISPR/Cas9-mediated knockout or knock-in cell lines

To generate Sec62-/- cells, human Sec62 targeting sequence (5′ AGTATCTTCGATTCAACTG 3′) was cloned into the gRNA expression vector (Mali et al., 2013) in order to direct Cas9 nuclease activity. HEK 293-Flp-In T-Rex cells were plated in a six-well plate and transfected at 70% confluence with 500 ng of the gRNA expression vector and 500 ng of the pSpCas9(BB)- 2A-Puro (Ran et al., 2013) plasmid using Lipofectamine 2000. Expression of Cas9 was selected by puromycin treatment (2.5 μg/ml) for 48 hr, after which cells were returned to non-selecting media for 72 hr. Cells were then plated at 0.5 cell/well in 96 well plates and expanded for 3 weeks. Individual clones were examined for Sec62 by immunoblotting. To engineer chromosomal 2X Strep-tagged Sec61α HEK293 cells, Sec61α gRNA sequence (5’ AAAGCGAGGTTGGCAGCATGGGG 3’) was cloned into the gRNA expression vector. The single-strand DNA oligonucleotide homology-directed repair (HDR) sequence (CATCTACCAGTACTTTGAGATCTTCGTTAAGGAGCAAAGCGAGGTTGGCAGCTCTGCCTGGAGCCACCCCCAGTTCGAGAAGGGCGGCGGCAGCGGCGGCGGCAGCGGCGGCGGCAGCTGGAGCCACCCCCAGTTCGAGAAGGCCTCTATGGGGGCCCTGCTCTTCTGAGCCCGTCTCCCGGACAGGTTGAGGAAGCTGC) was synthesized (IDT) for inserting 2X Strep-tag into the C-terminus of Sec61α encoding gene in HEK293 cells. 300 pmol of HDR oligonucleotide was electroporated into one million HEK293 cells along with 2.5 μg each of pSpCas9(BB)-2A-Puro and Sec61α gRNA plasmid (Amaxa kit R, program A-24; Lonza). Immediately after electroporation, cells were plated in a 6 well plate. After 24 h of electroporation, the expression of Cas9 was selected by puromycin treatment (2.5μg/ml) for 48 h. Cells were then returned to non-selecting media and grown for 2 days. Cells were replated at 0.5 cell/well in 96 well plates and expanded for 3 weeks. Individual clones were examined for the presence of 2X Strep-tagged Sec61α by immunoblotting with anti-Sec61α antibodies. Positive clones containing 2X Strep-tagged Sec61α exhibited a slower migrating band compared to Sec61α present in control HEK293 cells.

### Generation of stable cell lines

HEK 293 IRE1α-/- cells stably expressing IRE1α variants were previously described (Plumb et al., 2015). HEK293 Sec63-/- cells stably expressing Sec63 variants were created by transfecting with 1.8 μg of pOG44 vector and 0.2 μg of FRT vectors containing Sec63 using Lipofectamine 2000. After transfection, cells were plated in 150 μg/ml hygromycin and 10 ug/mL blasticidin. The medium was replaced every three days until colonies appeared. The colonies were picked and the protein expression was evaluated by immunoblotting.

### Immunoprecipitations

To test the interaction between IRE1α and the Sec61 translocon complex, 0.8 million HEK 293 cells were plated on a polylysine (0.1 mg/mL) coated 6 well plate. The cells were transiently transfected with 2 μg of HA-tagged or FLAG-tagged constructs using 5 μL of lipofectamine 2000 and treated with 100 ng/mL doxycycline unless otherwise indicated in the figure legends to induce protein expression. 24 h after transfection, cells were harvested in 1xPBS and centrifuged for 2 min at 10,000g. The cell pellet was lysed in 200 μL of Buffer A (50 mM Tris pH 8.0, 150 mM NaCl, 5 mM MgAc) including 2% digitonin by incubating on ice for 30 min. The 5% digitonin (EMD Millipore) stock was boiled for 5 min just before adding into Buffer A to avoid digitonin precipitating during IP. The supernatant was collected by centrifugation at 15,000*g* for 15 min. For co-immunoprecipitation, the supernatant was rotated with 12 μL of anti-FLAG-agarose (Biolegend) or 15 μL anti-HA magnetic beads (Thermo Scientific) for 1 h 30 min in the cold room. The beads were washed 3x with 1 mL of Buffer A including 0.1% digitonin. The bound material was eluted from the beads by directly boiling in 50 μL of 2x SDS sample buffer for 5 min and analyzed by immunoblotting.

To test the interaction between BiP and IRE1α, 0.8 million cells were plated on a 6 well plate and transiently transfected with 2 μg of IRE1α or its variants. Cells were washed and harvested in 1xPBS and centrifuged for 2 min at 10,000*g*. The cell pellet was lysed in either 200 μL of Buffer A including 2% digitonin, which preserves the IRE1α/Sec61/Sec63 complex, or NP40 buffer (50 mM Tris, pH 7.5, 150 mM NaCl, 0.5% deoxycholic acid, and 0.5% NP40), which disrupts the IRE1α/Sec61/Sec63 complex. Apyrase (10 U/mL) and 10 mM CaCl2 were included in both buffers and incubated for 30 min on ice. The cell lysate was centrifuged at 15,000*g* for 15 min. The supernatant was incubated with anti-HA magnetic beads (Thermo Scientific) for 1 h 30 min in the cold room. The beads were washed 3 times with either 1 mL of Buffer A including 0.1% digitonin or 1 mL of NP40 buffer and eluted by directly boiling in 50 μL of 2x SDS sample buffer for 5 min and analyzed by immunoblotting. HEK293 IRE1α-/- cells complemented with IRE1α-HA or IRE1α CNX-TMD-HA were plated as above and induced with 5 ng of Doxycycline for overnight. Cells were either treated or left untreated as indicated in figure legends and harvested, immunoprecipitated using digitonin buffer, and analyzed as above.

### Isolation of the endogenous IRE1α/Sec61/Sec61/BiP complex

To test the endogenous interaction between IRE1α and BiP, 7 million HEK 293 cells or 2X Strep-tagged Sec61α cells were plated on poly-lysine (0.1 mg/mL) coated 10 cm dishes. Second day, cells were harvested in 1xPBS and centrifuged at 1,500 g for 5 min. Cell pellets were lysed with 1 mL of lysis buffer (50 mM Tris pH 8.0, 250 mM NaCl, 5 mM MgAc, 2% digitonin) including 1x protein inhibitor, Apyrase (10 U/mL) and 10 mM CaCl2) for 30 min on ice. Cell lysates were centrifugated at 15,000 g for 15 min 4°C. The supernatant was incubated with either 20 μL anti-rabbit IRE1α antibodies (Santa Cruz) or control anti-rabbit His antibodies (Qiagen) for 1 h in the cold room. We then added 25 μL 50% Protein A slurry to each tube and rotated for another 1 h. For 2X Strep-tagged Sec61α pull down, the supernatant was directly incubated with 40 uL of 50% Strep-Tactin-XT beads (IBA, Germany) for 2 h in cold room. The beads were washed 3 times with 1 mL of wash buffer (50 mM Tris pH 8.0, 150 mM NaCl, 5 mM MgAc, 0.1% digitonin). The proteins were eluted from beads by directly boiling with 50 μL (for IRE1α immunoprecipitation) or 60 uL (for Strep pull down) of 2x SDS sample buffer for 5 min. The samples were analyzed by immunoblotting.

### Purification of the IRE1α complex and BiP

The IRE1α/Sec61/Sec63 complex and IRE1α Δ10 were purified as described previously (Sundaram et al., 2017). Briefly, microsomes were prepared from HEK293 cells stably expressing either 2xStrep IRE1α-FLAG or 2xStrep IRE1α Δ10-FLAG as described previously. 2 mL of microsomes (OD280 = 50) were lysed with an equal volume of lysis buffer (50 mM Tris pH8, 600 mM NaCl, 5 mM MgCl2, 2% digitonin (boiled prior to use) 1x protease inhibitor cocktail and 10% glycerol) by incubating 30 min on ice. The lysates were centrifuged at 25,000g for 25 min at 4°C. Supernatant was collected and passed through a column packed with 1 mL of compact StrepTactin beads (IBA, Germany) by gravity flow. Flow-through was collected and beads were washed with 6 x 1 mL of wash buffer (50 mM Tris pH 8.0, 150 mM NaCl, 5 mM MgCl2, 10% Glycerol and 0.1% digitonin). 2xStrep IRE1α or IRE1α Δ10-FLAG was eluted from the beads using 20 mM desthiobiotin (EMD Millipore) included in the wash buffer. The desthiobiotin eluted material was further purified by passing through a cation exchange chromatography (SP Sepharose beads, GE Healthcare). Briefly beads were prepared in a 2 mL Bio-Rad column and washed 5x using no salt buffer (20 mM Tris pH 6.0, 2 mM MgAc and 0.4% DBC). Purified protein was diluted 5x with no salt buffer and pass-through the S-column. Beads were washed 5x column volume with no salt buffer and eluted with 500 mM NaCl buffer (50 mM Tris pH8, 2 mM MgAc, 10% glycerol, and 0.4% DBC). BiP that is bound to IRE1α is mostly removed by this step because BiP does not efficiently bind to a cation exchange resin. Purified IRE1α/Sec63/Sec61 or IRE1α Δ10 were subjected to coomassie staining and quantified using BSA standards (Sigma).

The pET-28a (+) plasmid encoding N-terminally 6X His-tagged rat BiP lacking the N-terminal signal sequence was expressed and purified from E. coli as descripted by Amin-Wetzel et al., 2017. Briefly, pET-28a (+) His-BiP was transformed into BL21 Rosetta (DE3) cells. The overnight culture of His-BiP was innoculated into fresh liquid LB and grown to OD600 of ∼ 0.8 at 37°C. The culture was cooled down to 18°C and induce with 0.5 mM imidazole. After 16 h induction, the cells were harvested and resuspended with buffer A (50 mM Tris pH7.4, 500 mM NaCl, 10% glycerol, 1 mM MgCl2, 0.2% (v/v) Triton X-100, 20 mM imidazole). The suspension was passed through the high-pressure homogenizer for 4 times. The lysate was spun at 35000 rpm for 40 min at 4°C using Ti45 rotor. The supernatant was incubated with the prewashed 2 mL of Ni-NTA beads and washed with 20 mL of Buffer B (50 mM Tris pH 7.4, 500 mM NaCl, 10% glycerol, 1 mM MgCl2, 0.2% (v/v) Triton X-100, 30 mM imidazole). Subsequently, the column was washed with 10 mL of Buffer C (50 mM Tris pH 7.4, 1 M NaCl, 10% glycerol, 5 mM MgCl2, 1% (v/v) Triton X-100, 30 mM imidazole, 5 mM ATP) and further washed with 10 mL of Buffer D (50 mM Tris pH 7.4, 500 mM NaCl, 10% glycerol, 1 mM MgCl2, 30 mM imidazole).The bound proteins were eluted with Buffer E (50 mM Tris pH 7.4, 500 mM NaCl, 10% glycerol, 1 mM MgCl2, 250 mM imidazole). The peak fractions containing BiP was pooled and dialyzed against Buffer F (50 mM Tris pH 7.4, 150 mM NaCl, 10% glycerol, 5 mM MgCl2, 1 mM CaCl2). The purified proteins were flash frozen and stored at −80°C.

### In vitro reconstitution Sec61/Sec63-mediated BiP binding to IRE1α

IRE1α binding to BiP was adapted from Amin-Wetzel et al., 2017 with the following modifications. 12 μL of Anti-FLAG beads was incubated with either 0.15 μg of the 2X Strep-IRE1α-FLAG/Sec61/Sec63 complex or 0.15 μg of 2X Strep-IRE1α Δ10-FLAG in 500 uL of wash buffer (50 mM Tris pH 8.0, 150 mM NaCl, 10 mM MgCl2, 0.4% DBC) for 1 h at 4°C. The beads were washed twice with 1 mL of wash buffer. IRE1α bound beads were resuspended with 50 uL of binding buffer (50 mM Tris pH 8.0, 150 mM NaCl, 10 mM MgCl2, 1 mM CaCl2, 0.1% Triton X-100) including either BiP (1 μg) and ATP (2 mM). A negative control reaction was performed by incubating empty anti-FLAG beads with the buffer, BiP, and ATP. After incubation at 32°C for 30 min, the beads were quickly washed with ice-cold wash buffer including 2 mM ADP. The wash was repeated one more time with wash buffer excluding ADP. The bound proteins were eluted from beads 50 μL of 2X SDS sample buffer and analyzed by immunoblotting. We used the following protocol to BiP binding to IRE1α in Figure S6. 0.15 μg of the 2X Strep-IRE1α-FLAG/Sec61/Sec63 complex or 0.15 μg of 2X Strep-IRE1α Δ10-FLAG was incubated with and without 5 μg BiP in 50 uL of binding buffer (50 mM Tris pH 8.0, 100 mM NaCl, 10 mM MgCl2, 1 mM CaCl2, 2 mM ATP, 0.2% DBC) for 30 min at 32°C. A negative control reaction was performed by mixing the buffer, BiP, and ATP. The reactions were terminated by diluting with ice-cold NP40 buffer (50 mM Tris, pH 7.5, 150 mM NaCl, 0.5% deoxycholic acid, and 0.5% NP40) and incubated with 12 μL of anti-FLAG beads for 1 h 30 min at 4°C. After incubation, the beads were washed twice with 1 mL of NP40 buffer. The bound proteins were eluted by boiling beads with 50 μL of SDS sample buffer and analyzed by immunoblotting.

### Immunofluorescence

HEK293 IRE1α-/- cells stably complemented with IRE1α-HA (0.12 × 10^6^) were plated on 12 mm round glass coverslips (Fisher Scientific) coated with 0.1 mg/mL poly-lysine for 5 h in 24-well plates. For Figure 2A, the cells expressing IRE1α-HA were transfected with either 20 pmole of control or Sec62, or Sec63 siRNAs using 2 μL of lipofectamine 2000 and induced with 5 ng/mL of doxycycline to induce IRE1α expression. After 30 h of transfection, cells were treated with 5 μg/mL of thapsigargin (Tg) for 1.5 h before fixing and immunostaining as described previously (Sundaram et al., 2017). For Figure 2G, HEK293 IRE1α-/- cells stably expressing either WT IRE1α-HA or IRE1α CNX-TMD-HA were induced with 5 ng/mL doxycycline and treated with 5 μg/mL of Tm or Tg for the indicated time points. The treated cells were fixed and processed for immunostaining. For Figure 2D, the cells expressing IRE1α-HA were transfected with 0.1μg of Sec63 or Sec63 HPD/AAA using 1 μL of lipofectamine 2000. IRE1α expression was induced with doxycycline (5 ng/mL) for 16 h before treatment with 5 μg/mL Tg for 1.5 h followed by fixed and immunostained with anti-HA antibodies for IRE1α. The cells were imaged on Leica scanning confocal microscope and IRE1α clusters were quantified as previously described (Sundaram et al., 2017) with the following modifications. For each condition, we randomly chose at least 10 fields-of-view and took images. First, we identified the total number of cells per frame by manually counting Hoechst-stained nuclei. We counted more than 300 cells from the 10 images of each condition and looked for cells with IRE1α clusters. Of those cells, we calculated the percentage of cells with IRE1α clusters. Data was graphed using GraphPad Prism and represented with standard error of the mean (SEM) from two independent experiments.

### XBP1 mRNA splicing assay

Total RNA was extracted from cells using Trizol reagent (Ambion) according to the manufactures protocol. 2 μg of total RNA was treated with 1U/μL DNase I (Promega). 0.5 μg of DNAse-treated RNA was reverse transcribed into cDNA using Oligo(dT)20 primer (Qiagen) and M-MLUV reverse transcriptase (NEB). cDNA was amplified by standard PCR with TaqDNA polymerase using the primers: 5’-AAACAGAGTAGCAGCTCAGACTGC -3’, 5’-TCCTTCTGGGTAGACCTCTGGGAG -3’ (Calfon et al., 2002). PCR products of XBP1 were resolved by 2.5 % agarose gel and stained with ethidium bromide. The intensities of DNA bands were quantified on image analyzer (Image J, NIH).

### Phostag-based immunoblotting

Typically, 0.15 × 10^6^ cells were plated on 24 well poly-lysine coated plates. The following day, cells were treated with 2.5 μg/mL Tg for various time points indicated in Figure 3. The cells were directly harvested in 100 μL of 2X sample buffer and boiled for 5 to 10 minutes. IRE1α phosphorylation was detected by the previously described method (Yang et al., 2010). Briefly, 5% SDS PAGE gel was made using 25 µM Phos-tag (Wako). SDS-PAGE was run at 100 V for 2 h and 30 min. The gel was transferred to nitrocellulose (Bio-Rad, Hercules, CA) and followed with immunoblotting. The intensities of the Phos-tag bands were quantified on image analyzer (Image J, NIH). To probe the phosphorylation of PERK, the samples were run on a 7.5% Tris/Tricine gel for 2 h and 30 min and transferred to nitrocellulose membrane and blotted using a standard procedure.

### Quantification and statistical analysis

Quantification of immunoblots were performed using image J gel analysis/ lane plotting. Error bars represent standard error of the mean (SEM) from two or three independent experiments as indicated in the figure legends. *p*-values were calculated by the GraphPad Prism using Student’s *t* test. *p*-values < 0.05 was considered as significant.

